# Chromatin regulator HELLS mediates SSB repair and responses to DNA alkylation damage

**DOI:** 10.1101/2024.12.19.629292

**Authors:** Joyous T Joseph, Christine M Wright, Estanislao Peixoto, Etsuko Shibata, Asad Khan, Yong Li, Jason S Romero Neidigk, Olivia Decker, Krishna Reethika Kadali, Azait Imtiaz, Brianna A Jones, Yangfeng Zhang, Sergio A Gradilone, Zachary A Lewis, Rafael Contreras-Galindo, Arko Sen, Anindya Dutta, Wioletta Czaja

## Abstract

The SNF2 family chromatin remodeler HELLS has emerged as an important regulator of cell proliferation, genome stability, and several cancer pathways. Significant upregulation of HELLS has been reported in 33 human cancer types. While HELLS has been implicated in DNA damage response, its function in DNA repair is poorly understood. Here we report a new regulatory link between HELLS and single-strand break (SSB) repair in cellular responses to DNA alkylation damage. We found that loss of HELLS impairs SSB repair, and selectively sensitizes cells to DNA alkylating agents and PARP inhibitors (PARPi). Furthermore, we found that HELLS is co-expressed with PARP1 in cancer cells, and its loss is synthetic lethal with homologous recombination deficiency (HRD). This work unveils new functions of HELLS in modulating SSB repair and responses to clinically relevant DNA alkylation damage, thus offering new insights into the potential therapeutic value of targeting HELLS in cancer.

**Graphical Abstract:** 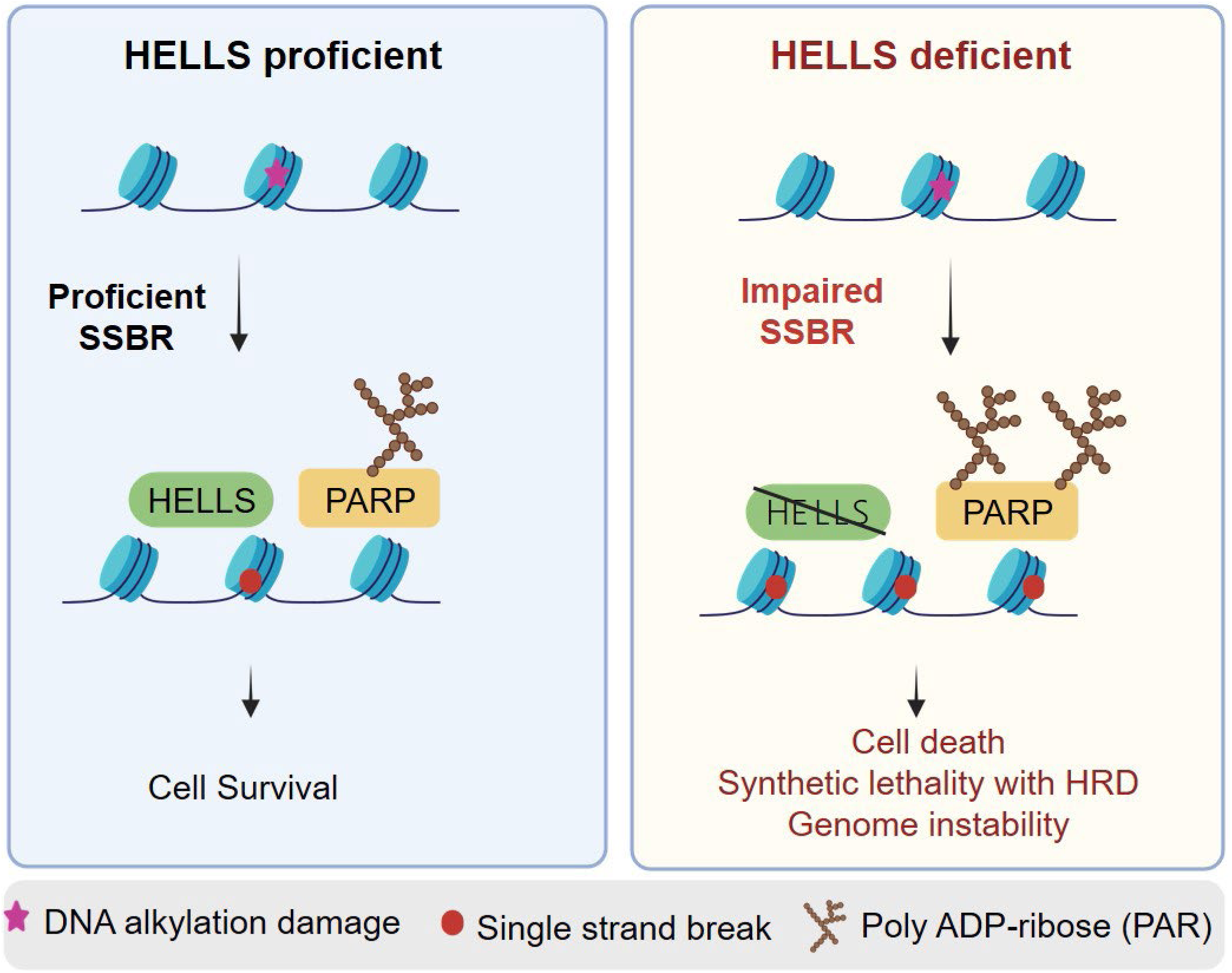

## Introduction

Proficient and tightly regulated DNA repair is critical for human health. Defective DNA repair is associated with various disorders including immunodeficiency, developmental abnormalities, neurodegeneration, and cancer [1, 2]. In eukaryotes, DNA is packaged into chromatin, which undergoes dynamic remodeling in response to DNA damage [3]. The SNF2 family of ATP-dependent chromatin remodeling (ACR) enzymes mediate chromatin reorganization and play crucial roles in regulating the DNA transcription and repair processes in the context of chromatin [4]. Several distinct SNF2 remodelers have been implicated in regulation of DNA damage response (DDR) and cancer pathways in multiple cancer types, highlighting their potential utility as drug targets [5–8].

DNA alkylating agents are frequently employed in cancer chemotherapy, inducing the formation of DNA base damage, cytotoxic SSB and DSB damage in DNA. Various DNA repair pathways mediate responses to DNA alkylation, mainly Base Excision Repair (BER) and Homologous Recombination (HR) [9–12]. Highly proliferating cancer cells are particularly susceptible to alkylating agent-induced DNA damage and they must heavily rely on proficient BER and HR for cell survival. Therefore, deficiencies in BER and/or HR can potentiate cancer cell sensitivity to DNA alkylating agents [11, 13, 14].

BER is the frontline DNA repair pathway mediating the resolution of DNA alkylation damage. BER involves the action of specific glycosylases and endonucleases, including MPG and APE enzymes, that recognize and excise damaged DNA bases, leading to the formation of abasic sites (AP sites) and single-strand breaks (SSB) [15, 16]. Packaging of DNA in nucleosomes presents a barrier to efficient SSB repair [15]. Indeed, it has been demonstrated that POLβ catalyzed insertion of nucleotide during SSB repair is strongly inhibited by nucleosomal core particle at the basic level of chromatin organization [17]. PARP1 (Poly(ADP-ribose) polymerase 1) plays a crucial role in the repair of single-strand breaks (SSB) in the context of chromatin. Upon binding to SSB, PARP1 becomes activated and catalyzes the addition of poly(ADP-ribose) (PAR) chains onto itself and other target proteins, a process known as PARylation. PARP1-mediated PARylation helps to relax chromatin structure, and facilitates the recruitment of several SNF2 enzymes, including ALC1, CHD3, and ISWI, further promoting the recruitment and accessibility of BER repair factors, including XRCC1, POLβ, and LIG3 to the damaged DNA [18]. The XRCC1 promotes PARP1-dependent recruitment of PARP2 to damage sites [19].The unrepaired and persistent SSBs and AP sites are highly cytotoxic BER intermediates that can block the progression of DNA replication forks and lead to the generation of S-phase-dependent DSB [20]. Specifically, persistent SSBs can cause DNA replication fork collapse and replisome disassembly [21]. Abundant, unrepaired SSB can lead to hyperactivation of PARP1 and increased PARP1 retention on damaged DNA, known as “PARP trapping”. Similarly, PARP inhibitors (PARPi) can lead to excessive accumulation and trapping of PARP on the DNA, creating highly cytotoxic PARP-DNA complexes [22–25]. Removal of trapped PARP from chromatin is mediated by several mechanisms involving the activity of SNF2 family remodeler, ALC1 [26, 27] and recently reported p97-mediated pathway [28]. In addition, XRCC1 prevents toxic PARP1 trapping during BER [29].

The HELLS (helicase, lymphoid specific; also known as SMRCA6, LSH, PASG) protein is a member of the conserved SNF2 family of ATP-dependent chromatin remodelers [4, 30]. Mutations in *HELLS* cause immunodeficiency-centromeric instability-facial anomalies (ICF4) syndrome [31]. Although *HELLS-/-* mice die prenatally, mutational analysis of the *HELLS* gene in ICF4 patients suggests that loss of HELLS is compatible with human life [31]. HELLS is expressed in highly proliferating cells of the lymphoid tissue, germ cells, stem cells, and it is substantially elevated in cancer cells [30]. HELLS has been implicated in several cancer pathways, including proliferative signaling, genome instability, deregulated cell energetics, and invasion, all of which influence many aspects of cancer initiation, progression, and responses to therapy [7]. Significant upregulation of HELLS has been reported in 33 human cancer types, including breast, leukemia, and glioblastoma [32]. The clinical significance of HELLS in glioblastoma is highlighted by the poor prognosis of glioblastoma patients with elevated HELLS expression and the improved survival of glioblastoma mouse models upon HELLS downregulation [33]. HELLS has been proposed to serve as a potential biomarker for cancer diagnosis and prognosis and to offer value in immune-based, targeted, or cytotoxic therapies [32, 34]. Cells deficient in HELLS are characterized by decreased proliferation, increased levels of genomic instability, senescence, and sensitivity to genotoxic agents, the phenotypes frequently associated with deficient DNA repair pathways [7]. Human HELLS has been implicated in the DNA damage response (DDR) and HR-mediated repair of gamma radiation-induced DSB at heterochromatin [35, 36]. Another study linked HELLS with regulation of the classical non-homologous end joining (c-NHEJ) pathway [37]. A recent study by Xu *et al*. found that HELLS plays a crucial role in maintaining replication fork stability in response to replication stress by promoting the deposition of the macroH2A histone variant, RAD51 filament formation and protecting stalled forks from nucleolytic degradation [38].

Here, we report a new regulatory link between the HELLS remodeler, SSB repair, and cellular responses to DNA alkylation damage. We found that HELLS-deficient cells exhibit enhanced selective sensitivity to DNA alkylating agents, transient increase in unrepaired SSBs associated with deficient association of BER proteins with damaged chromatin, hyperactivation of PARP1, elevated PAR levels, progressive accumulation of alkylation-derived DSB, G2/M cell-cycle arrest and induction of apoptosis. Furthermore, we found that HELLS deficiency confers PARPi sensitivity, sensitizes BRCA2 downregulated cells to DNA alkylation damage, and results in synthetic lethality when combined with RAD51 inhibition. The alkylation sensitivity, and synthetic lethality phenotypes are consistent with deficient SSB repair that can lead to the formation of S-phase-dependent DNA damage creating a strong dependency on the functional HR pathway for cell survival. Our studies identify HELLS as an important regulator of the BER/SSBR pathway, that could serve as a potential therapeutic target in the sensitization of cancer cells to alkylation chemotherapy and PARPi, especially in the context of HR deficiency.

## Results

### Loss of HELLS leads to decreased cancer cell proliferation and increased cell type-specific selective sensitivity to DNA alkylating agents

HELLS protein expression is significantly elevated in various malignant cell lines, including those derived from myeloid leukemia [30]. To investigate the role of HELLS in DNA damage repair, we utilized human cancer cell lines with high HELLS expression level, including myeloid leukemia HAP1 cells, and cervical cancer HeLa cells. A non-cancer, immortalized human fibroblast CHON-002 cells expressing very low levels of HELLS protein were also used. The *HELLS* gene was CRISPR edited to create a single nucleotide change resulting in the complete loss of HELLS protein (HELLS KO) in HAP1, HeLa, and CHON-002 backgrounds. Both the HAP1 and HeLa parental strains exhibit abundant endogenous HELLS expression, consistent with high proliferation rates. (**Fig. 1A and Fig. 2A**). The non-cancerous, slowly proliferating, CHON-002 fibroblasts display very low levels of HELLS protein (**Fig. 2A**).

**Figure 1.**
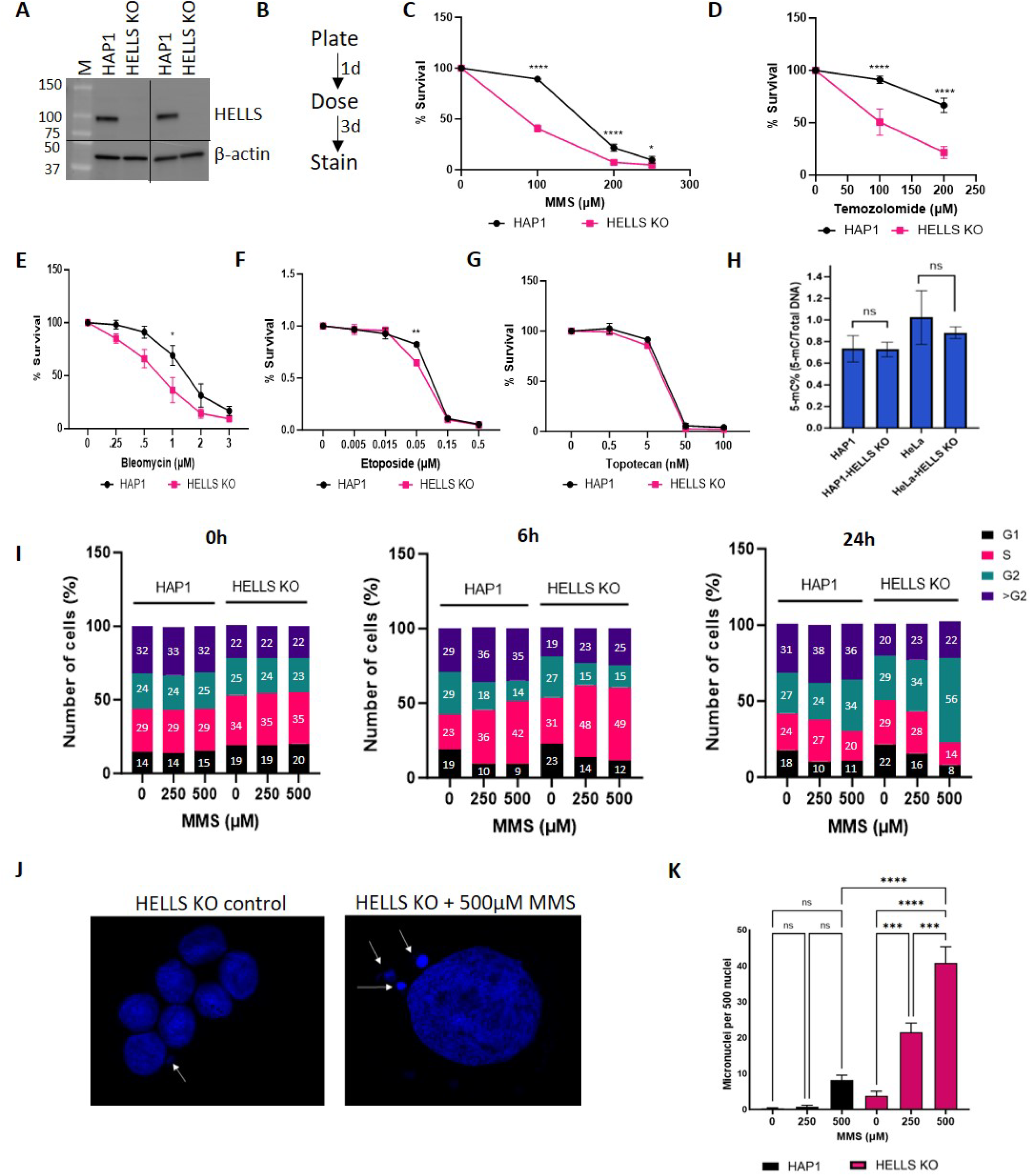
HELLS promotes cancer cell survival and genome stability in response to DNA alkylation damage. **A**. Western blot of Whole Cell Extract (WCE) in HAP1 and HELLS KO probed for HELLS. Actin was used as a loading control. **B**. Schematic representation of survival assays using crystal violet staining. **C-F**. Quantification of crystal violet cell survival for **C**. MMS **D**. Temozolomide **E**. Bleomycin or **F**. Etoposide. Data are mean ± SEM. n=3 independent biological replicates. Two-way ANOVA with Tukey’s multiple comparisons was used to determine significance. **G**. HAP and HELLS KO sensitivity to Topotecan was determined by cck-8 after treatment to the indicated doses for 48hr. Data are mean ± SEM. n=3 independent biological replicates. **H**. Quantification of 5-methyl cytosine (5-mC) content of parental and HELLS KO cell of HAP1 and HeLa using the MethylFlash^TM^ Global DNA Methylation ELISA kit. n=4 independent biological replicates. Statistically significant differences were calculated using unpaired t-test. **I.** Relative cell cycle distribution of cells after treatment with MMS. n=3 independent biological experiments. **J.** Representative image of micronuclei in HELLS KO cells. **K.** Formation of micronuclei in response to MMS treatment. Micronuclei formation in HELLS KO HAP cells treated with MMS. Cells were treated with 250μM or 500μM MMS for 1hr. The number of micronuclei was visualized by DAPI staining in HAP1 control or HELLS KO HAP1 cells 48 hr post MMS treatment. Micronuclei counts were performed in 4 microscopy fields in each experimental setting. Statistically significant differences were calculated using one-way ANOVA with Tukey’s multiple comparisons. Data are mean ± SEM and are representative of at least two independent experiments. (*p<0.05, **p<0.01, ***p<0.001, ****p<0.0001) with DNA methylation changes.

**Figure 2.**
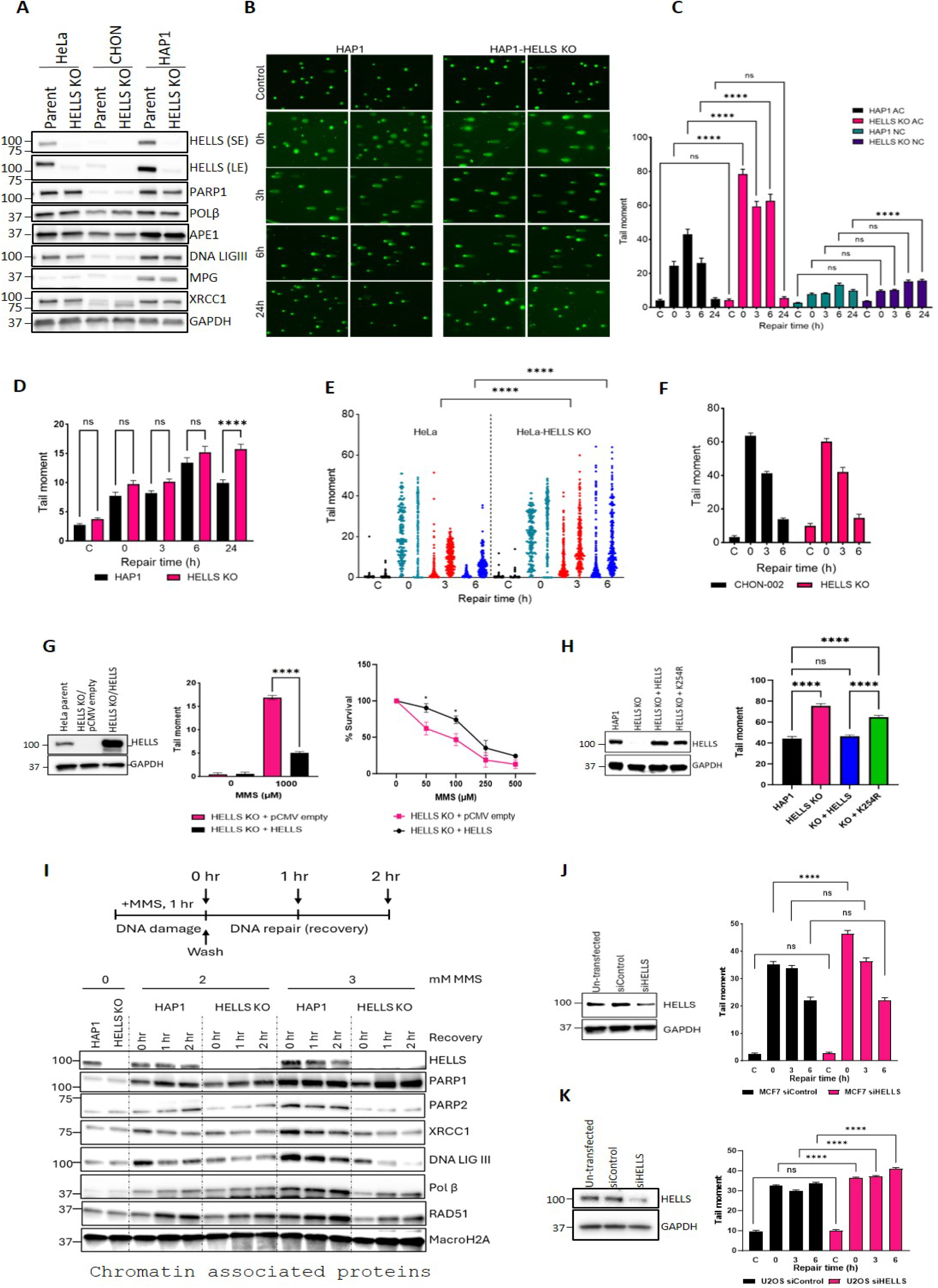
Loss of HELLS results in deficient repair of alkylation-derived DNA Single single-strand breaks (SSBs) in specific cell types. **A**. HELLS and key BER DNA repair protein expression levels in 3 cell lines as visualized by Western blot in WCE. SE=short exposure LE=long exposure. GAPDH was used as a loading control. **B**. Alkaline comet (AC) images of HAP1 and HAP1-HELLS KO cells following treatment with 500μM MMS for 1hr. **C**. DNA strand breaks quantified by alkaline (AC) and neutral (NC) assays. Data are the average of comet tail moments of 50 cells per sample. n=3 independent biological experiments. **D**. Double strand breaks quantified based on NC assay in HAP1 cells. Graph is plotted as in **C**. **E**. DNA strand breaks quantified by AC assay in HeLa cells following treatment with 1mM MMS for 1hr. Data represent individual comet tail moments of 200 cells per sample per experiment plotted vertically. Two independent experiments are displayed side by side. **F**. DNA strand breaks quantified by AC assay in CHON-002 cells at indicated time points following treatment with 500μM MMS for 1hr. Graph is plotted using the average of comet tail moments of 100 cells per sample per experiment of two independent experiments. **G**. Western blot of WCE in HeLa-HELLS KO cells transiently transfected with pCMV empty or pCMV-HELLS (HELLS overexpression) vectors, probed for HELLS. GAPDH was used as a loading control. DNA strand breaks quantified by AC assay in HeLa HELLS KO cells transfected with pCMV-HELLS following treatment with 1mM MMS for 1hr. Data represent mean tail moments of 200 cells per sample. n=2 independent biological replicates. HeLa HELLS KO cells transiently transfected with pCMV-empty vector and pCMV-HELLS were exposed to MMS for 48hr and cell survival was determined with CCK8 reagent. Data are mean ± SEM. n=3 independent biological replicates. **H**. Western blot of WCE in HAP1 HELLS KO cells containing the stable genomic integration of HELLS WT, and HELLS K254R (catalytically inactive). GAPDH was used as a loading control. DNA strand breaks quantified by AC assay in the cell lines. Data are the average of comet tail moments of 100 cells per sample of two independent biological experiments. **I**. HAP1 and HAP1-HELLS KO cells were treated with the indicated doses of MMS for 1hr. The media was changed, and cells were allowed to recover for 0hr, 1hr, or 2hr. BER pathway members were visualized on chromatin by Western blot. MacroH2A was utilized as a loading control. **J**. Western blot of WCE in MCF7 cells probed for HELLS indicating downregulation of HELLS by siRNA for 72h. GAPDH was used as a loading control. DNA strand breaks quantified by AC assay in MCF7 cells followed by HELLS downregulation. Cells transfected with 3nM siRNA for 72h were dosed with 1mM MMS for 1 hr. Data is the average of comet tail moments of 100 cells per sample. **K**. Western blot of WCE in U2OS cells probed for HELLS indicating downregulation of HELLS by siRNA for 48h. GAPDH was used as a loading control. DNA strand breaks quantified by AC assay in U2OS cells followed by HELLS downregulation. Cells transfected with 3nM siRNA for 48h were dosed with 1mM MMS for 1 hr. Data is the average of comet tail moments of at least 200 cells per sample. n=2 independent biological experiments. Statistical significance for all comet assays was determined by two-way ANOVA of the mean tail moments with Tukey’s multiple comparisons test. (*p<0.05, **p<0.01, ***p<0.001, ****p<0.0001).

Under normal growth conditions, HELLS-deficient cells in the HAP1 and HeLa, but not CHON-002 cells, demonstrated reduced proliferation rates as compared with the parental counterparts (**Fig. S1A**). Decreased proliferative capacity in HELLS KO cells is consistent with other studies [33, 37, 39] and highlights an important function of HELLS in cancer cell growth and proliferation.

To determine the impact of HELLS loss on cell survival in response to DNA damage, cells were exposed to DNA alkylating agents, including Methyl Methanesulfonate (MMS) and Temozolomide (TMZ), as well as other chemotherapeutics such as Bleomycin (BLM), Etoposide (ETP) and Topotecan (TPT) (**Fig. 1B-G**). HELLS-deficient HAP1 cells displayed significant sensitivity in response to MMS and TMZ (**Fig. 1C-D),** but very mild, or lack of sensitivity in response to BLM, ETP and TPT (**Fig. 1E-G**). Notably, we found that HELLS-deficient HeLa and CHON-002 cells, unlike HAP1 HELLS KO cells, did not exhibit increased sensitivity to MMS alone (**Fig. S1B&E**), suggesting the existence of compensatory mechanisms that suppress the alkylation sensitivity phenotype in certain cellular backgrounds. These data underscore the cell type-specific functions of HELLS in mediating cell survival in response to DNA alkylation, and are consistent with the context-specific functions of SNF2 remodelers [4]. Importantly, our results suggest selective sensitivity of HELLS-deficient cells to DNA alkylators, and recapitulate MMS sensitivity phenotypes reported in HELLS-deficient plant and fungal cells, highlighting the evolutionarily conserved role of HELLS in response to DNA alkylation damage [40–42].

Several studies reported that loss of HELLS in mouse and human cells leads to decreased global DNA methylation [43–45]. To determine if loss of HELLS impacts global CpG methylation levels, we have performed CpG ELISA, and found that loss of HELLS does not impact levels of CpG methylation in HAP1 cells. Similarly, we detected slightly decreased level of CpG methylation in HELLS KO HeLa cells as compared with the parental cells, however the differences were not statistically significant. (**Fig 1.H**). This data suggests that the alkylation sensitivity phenotypes observed in HELLS KO cells are mediated by the loss of HELLS, and are likely not associated

The haploid HAP1 cell line is a suitable model to study genetic interactions between DNA repair pathways [46–48]. The near haploid HAP1 cells are inherently unstable and tend to spontaneously transition to the diploid state during prolonged passage in cell culture [49]. Ploidy status might influence DNA repair pathway choice and overall sensitivity to DNA damage [50]. Therefore, we aimed to determine whether the ploidy state in HAP1 affects the MMS sensitivity phenotypes in HELLS-deficient cells. To achieve this, HAP1 cells were single-cell sorted via FACS to obtain pure populations of haploid and diploid cells, which were subsequently tested for MMS sensitivity. We observed that ploidy status did not significantly impact sensitivity to DNA alkylation. Both haploid and diploid HELLS KO clones exhibited comparable levels of sensitivity and cell survival in response to MMS treatment (**Fig. S2A**). The ploidy status of these clones and their HELLS expression level were verified by flow cytometry and western blotting (**Fig. S2B&C**).

#### Loss of HELLS results in G2/M cell cycle arrest and elevated micronuclei in response to DNA alkylation damage

To determine the impact of HELLS deficiency on the cell cycle, unsynchronized cells were exposed to increasing doses of MMS for 1hr. After MMS removal, the cells were incubated in drug-free media for 6 and 24hr, and cell cycle progression was assessed by flow cytometry (**Fig. 1I**). We found that untreated HELLS KO cells had a slightly increased population of cells in S-phase (34%) compared to the parental cell line (29%). MMS-induced DNA alkylation damage is known to block S-phase progression, leading to transient accumulation of cells in S-phase [51]. As expected, exposure to 500µM MMS resulted in a distinct increase in the S-phase population at 6hr post-MMS treatment: 42% in HAP1 cells and 48% in HELLS KO cells (**Fig. 1I**).

Persistent, unrepaired DNA alkylation damage is known to induce G2/M cell cycle arrest and cell death which prevents genome instability in subsequent cellular generations [51]. A prominent G2/M cell cycle arrest was observed in HELLS-deficient cells 24hr post-MMS exposure, with 55% of cells arrested in G2/M, compared to 34% in HAP1 cells. This data is consistent with findings in other cells defective in DNA repair, especially SSB repair, which also exhibit G2/M cell cycle arrest in response to MMS-induced DNA damage [52]. Unrepaired or miss-repaired DNA damage can lead to chromosome breaks and segregation defects, which commonly contribute to the formation of micronuclei [53]. Therefore, we sought to determine whether loss of HELLS results in elevated micronuclei in response to DNA alkylation damage. Cells were exposed to 250 and 500µM MMS for 1hr, followed by drug removal and cell recovery in fresh media for 48hr. Micronuclei formation was assessed 48hr post-MMS exposure. We observed an MMS dose-dependent increase in micronuclei formation that was substantially elevated in HELLS-deficient cells. In HAP1 cells, treatment with 500µM MMS led to ∼10 micronuclei per 500 nuclei. Whereas, in HELLS-deficient cells, this dramatically increased to ∼40 micronuclei per 500 nuclei and a significant increase was also detectable at lower MMS concentrations (**Fig. 1J&K**). Elevated micronuclei and G2/M cell cycle arrest have also been detected in XRCC1 mutant cells deficient in SSB repair in response to MMS [52]. Our findings are consistent with earlier studies reporting elevated micronuclei in HELLS-deficient cells [36] and further highlight the impact of DNA alkylation damage on micronuclei formation in HELLS-deficient cells.

### Loss of HELLS results in deficient repair of alkylation-derived Single Strand Breaks (SSB)

Given the prominent and selective sensitivity of HAP1 HELLS deficient cells to DNA alkylating agents, associated with G2/M arrest and elevated micronuclei, we aimed to explore the involvement of HELLS in the repair of DNA alkylation damage and modulation of the BER pathway. We found that loss of HELLS did not affect endogenous levels of the key BER proteins, including PARP1, POLβ, APEX1, DNA LIGIII, MPG, and XRCC1 (**Fig. 2A**). Notably, the non-tumor CHON-002 cells expressed substantially less BER proteins than either HAP1 or HeLa cancer cells, irrespective of the HELLS status. This is consistent with the assumption that non-cancerous cells may have low endogenous DNA damage, and therefore have lower levels of DNA repair proteins, as compared to cancerous cells with elevated endogenous DNA damage and increased levels of DNA repair proteins [54]. HeLa cells also display substantially lower MPG levels than HAP1 cells **(Fig. 2A**), implying inherent differences in the status of the BER pathway in diverse cell lines.

To determine the contribution of HELLS to SSB repair, cells were exposed to 500µM MMS for 1hr, followed by MMS removal and repair in fresh, drug-free media for 3, 6, and 24hr. The formation and repair of MMS-induced alkylation-derived SSB and DSB were assessed using alkaline (AC) and neutral comet (NC) assays. Under normal growth conditions HELLS deficient cells (unexposed to MMS), did not display an increase in the comet tail moment, and were comparable with their parental counterparts, as evident in both AC and NC assays (**Fig. 2C-E**). This finding is consistent with previous studies showing that XRCC1 mutant cells, which are deficient in SSB repair, do not exhibit detectable endogenous DNA damage under normal growth conditions, as assessed by alkaline comet [29]. Notably, we found that HELLS-deficient HAP1 and HeLa cells accumulate high levels of SSBs in response to MMS, which persist for 3- and 6-hr post-MMS removal (**Fig. 2C&E**) as demonstrated by AC comet assays. We also detected increased SSB is siRNA HELLS downregulated MCF7 and U2OS (**Fig.2J&K**). In contrast, non-tumor HELLS-deficient CHON-002 cells appeared to repair SSB proficiently (**Fig. 2F**). This data suggests that HELLS modulates SSB repair in specific cell types. We speculate that highly proliferating tumor cells might specifically rely on HELLS for proficient SSB repair, while HELLS might be dispensable for SSB repair in slowly proliferating, non-tumor cells.

It has been suggested that MMS does not directly induce abundant DSB [55]. However, one-ended DSB can arise if replication forks collide with unrepaired and persistent BER repair intermediates, such as SSB [56]. To assess the contribution of HELLS to the formation and repair of alkylation-derived DSB, we performed neutral (NC) comet assays (**Fig. 2D**). As expected, MMS-induced DSBs arise progressively over time, however at much lower frequencies as compared with high levels of SSBs (DSB tail moment: ∼5-20, versus SSB tail moment: ∼20-80) (**Fig. 2C&D**). Low levels of alkylation-derived DSB were detected at the early repair time points and were progressively increasing at similar rates in both HELLS-proficient and HELLS-deficient HAP1 cells between 1hr and 6hr post-MMS exposure, suggesting that loss of HELLS does not affect DSB repair under these conditions. Significantly elevated DSBs in HELLS KO cells at 24hr post-MMS likely represent a fraction of SSB- derived DSBs arising during S-phase. The neutral comet studies were complemented with the immunofluorescence studies monitoring γH2AX, a well-established marker of DSB. We observed that alkylation-induced γH2AX foci were substantially elevated in HELLS KO cells and persisted for extended periods of 48- and 72-hr post-MMS exposure (**Fig. S3A&B.**) The elevated γH2AX likely represents a mix of alkylation-derived DSB formed at replication forks that collapsed at unrepaired SSB, as well as DSB generated in the early stages of apoptosis. Indeed, MMS-treated cells showed increased levels of apoptosis detected by annexin staining and flow cytometry analysis (**Fig. S3C**). The HELLS KO cells are unlikely to be deficient in homologous recombination (HR), as they do not display increased sensitivity to the DSB-inducing agent ETP and TOP (**Fig. 1F&G**). Furthermore, at least three independent studies, using the HR-GFP reporter assay in MCF10A, U2OS cells, and HEK293T concluded that HELLS deficiency does not compromise the capacity of the canonical HR pathway in these cell types [36, 38, 57]. However, since HELLS has been reported to promote DSB repair in the heterochromatin, it is possible that the repair of alkylation-derived DSB arising in heterochromatin might be deficient in HELLS KO cells [36].

To investigate whether HELLS overexpression impacts rates of SSB repair and sensitivity to MMS, HeLa HELLS-deficient cells were transfected with pCMV plasmid driving overexpression of HELLS, or pCMV empty vector. Overexpression of HELLS protein in HELLS KO cells contributed to a rapid decrease of SSBs within 1hr of MMS exposure, consistent with robust repair (**Fig. 2G**). HELLS overexpression also contributed to increased MMS resistance in cell survival assay (**Fig. 2G**). To determine if the ATPase activity of HELLS is required for the repair of SSBs, we utilized plasmid constructs expressing WT HELLS, and catalytically inactive mutant HELLS, containing a K254R substitution in the Walker A box of ATPase domain (**Fig. 2H**). The K254R mutation prevents HELLS protein from binding ATP and likely disables its ATP-dependent chromatin remodeling, as reported in a previous studies [36]. WT HELLS construct and catalytically inactive K254R HELLS construct were stably integrated into the HAP1-HELLS KO cells, and their level of expression was comparable with endogenous HELLS expression in the parental HAP1 cell line (**Fig. 2H**). We found that catalytically inactive HELLS mutant displayed significantly elevated SSBs in the alkaline comet assays implying that chromatin remodeling activity of HELLS is important for SSB repair (**Fig. 2H**). Interestingly, we found that both WT and catalytically inactive HELLS rescued MMS sensitivity in HELLS KO cells, suggesting that chromatin remodeling activity of HELLS is not essential to promote cell survival in response to MMS (**Fig.S4A**). These data suggest that proficient SSB repair relies on HELLS ATP-dependent catalytic activity, but its catalytic activity appears to be dispensable in promoting cell survival in response to DNA alkylation damage.

#### Loss of HELLS impairs the recruitment of DNA repair proteins to damaged chromatin

To determine the contribution of HELLS to the recruitment of BER proteins to chromatin in response to DNA alkylation damage, we performed chromatin fractionation experiments. Cells were exposed to 2 and 3mM MMS for 1hr, followed by drug removal and cell recovery during a period of 1 and 2hr. Chromatin fractions were isolated, and chromatin-associated proteins were detected by western blotting (**Fig. 2I**). In order to detect changes in chromatin-associated proteins in response to MMS, milimolar concentrations of MMS were used following previously published protocols [26, 58]. Under these conditions, both HAP1 and HELLS KO cells retained cellular morphology and no signs of cell shrinking, or death were detected during the recovery period of 1-2hr. We found that in the absence of external DNA damage loss of HELLS protein does not impact endogenous levels of chromatin-associated BER proteins (**Fig. 2I**). In response to increasing dosage of MMS, we detected abundant recruitment of PARP1 to chromatin, in both HELLS proficient and deficient cells, suggesting that HELLS does not impact PARP1 enrichment on chromatin in response to MMS (**Fig. 2I**). In addition, levels of MPG, APE1, PARG1, and XRCC5 were not altered in HELLS KO cells (**Fig. S6**). Notably, we detected lower enrichment of XRCC1, and XRCC1-associated proteins POLβ, and LIG3 in HELLS deficient cells in response to increasing concentrations of MMS (**Fig.2I**). Furthermore, we found that PARP2 enrichment on chromatin was also reduced in HELLS KO cells. Recent studies demonstrated that XRCC1 acts upstream of PARP2, and it mediates enrichment of PARP2 to DNA damage sites [19], therefore decreased XRCC1 enrichment in HELLS KO cells likely explains lower PARP2 levels on chromatin in response to MMS-induced DNA damage. PARP2 has been implicated in stabilizing replication forks in response to DNA alkylation damage through regulation of RAD51 [59]. Furthermore, RAD51 has been shown to promote replication through alkylated DNA damage, mediating fork protection, independently of its role in DSB repair [59–61]. We found that RAD51 accumulation on chromatin was decreased in HELLS KO cells in response to MMS. These data demonstrate that MMS-induced chromatin enrichment of several key BER enzymes as well as RAD51 is impaired in the absence of HELLS, suggesting that in addition to deficient SSBR, the fork protection mechanisms could be also deficient in HELLS KO cells.

### Loss of HELLS results in sensitivity to Olaparib, and synergistic hypersensitivity when Olaparib is combined with non-toxic doses of MMS

Previous studies have reported that SSB-deficient cells display increased sensitivity to PARP inhibitors [14]. Inhibition of PARP activity leads to the accumulation of persistent SSB and to the generation of lethal DSB at collapsed replication forks during S-phase, when cell proliferates [62, 63]. Since HELLS KO cells are deficient in SSB we hypothesized that these cells will be more sensitive to PARP inhibitors. Indeed, we found that HAP1 HELLS KO cells treated with Olaparib (OLA) exhibit decreased cell viability (**Fig. 3A&B**). Our data is consistent with several large-scale CRISPR screen-based studies that also have identified HELLS as a key mediator of PARPi sensitivity in various cell lines, including those from triple-negative breast cancer, ovarian cancer, and prostate cancer [64–66].

**Figure 3.**
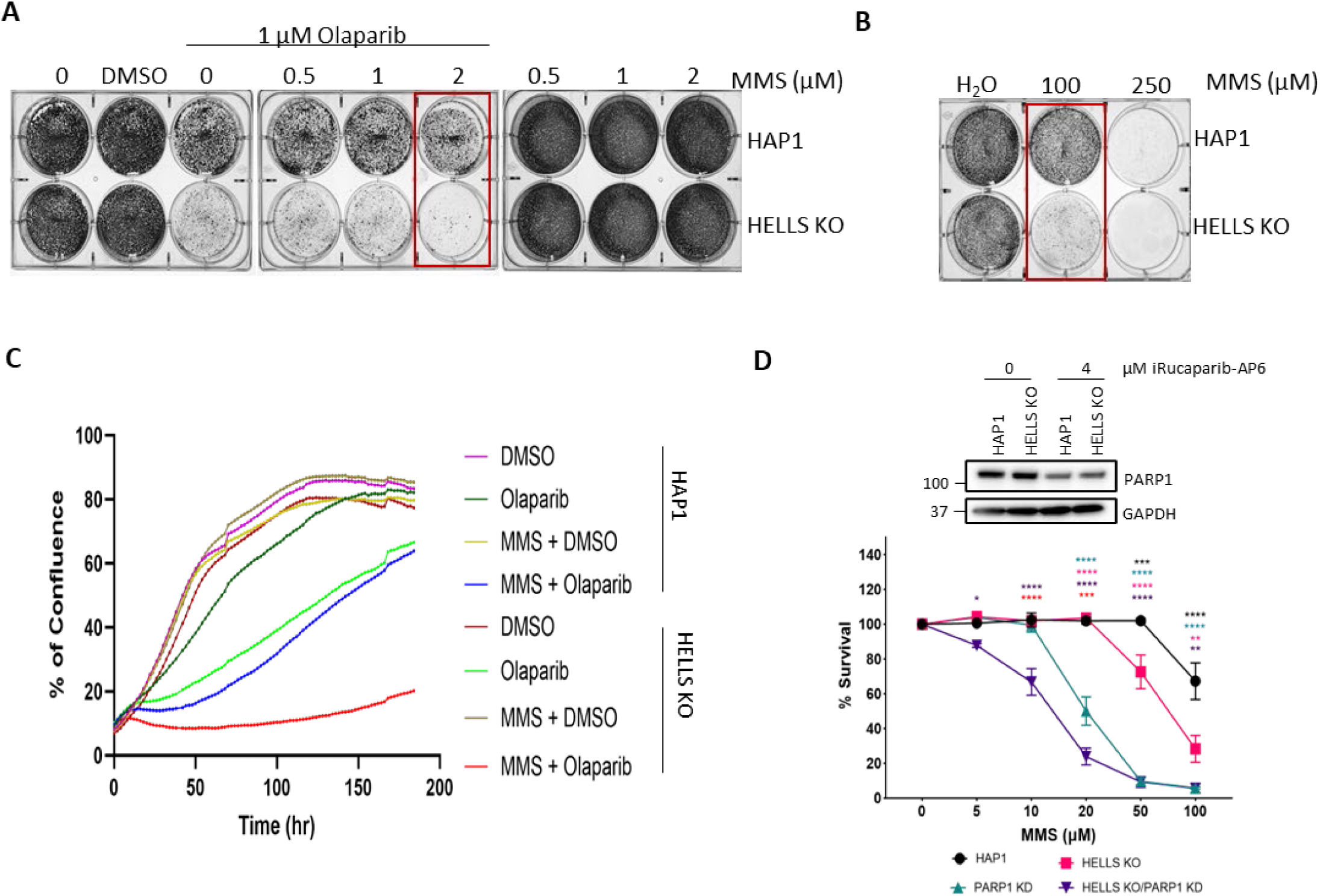
HAP1 cells lacking HELLS display hypersensitivity to MMS combined with Olaparib. **A.** Representative image of cell survival assay of HAP1 and HAP1-HELLS KO cells treated with indicated concentrations of MMS and Olaparib for 3d. Media was changed, and cells were grown for 2 additional days. Data represents at least 4 independent experiments. **B.** Cell survival assay of HAP1 and HAP1-HELLS KO cells treated with indicated concentrations of MMS 4d. Data represents at least 4 independent experiments **C.** Proliferation curves of HAP1 and HELLS KO cells are presented as the percentage of confluence for cells treated for 1hr with 20µM MMS, 1µM Olaparib, or their combination. Data is a representation of at least two independent experiments. **D.** Cell survival followed by PARP1 degradation assessed by CCK8. HAP1 and HELLS KO cells were exposed to 4μM iRucaparib-AP6 and indicated doses of MMS for 48hr to induce PARP1 degradation and DNA alkylation damage. Asterisk colors denote the following comparisons: black-HAP1 to HELLS KO; cyan-HAP1 to PARP1 KD; pink-HELLS KO and PARP1 KD; purple-HELLS KO and HELLS KO/PARP1 KD; red-PARP KD and HELLS/PARP KD. Western blot of WCE in HAP1 and HELLS KO cells probed for PARP1, indicating PARP1 degradation by iRucaparib-AP6.

Furthermore, PARP inhibitors are known to potentiate the cytotoxicity of DNA alkylating agents by inhibiting SSB repair, and inducing the formation of highly cytotoxic PARP-DNA-SSB (trapped PARP) adducts as well as single-stranded DNA gaps at replication forks [22, 67]. Indeed, we found that HAP1 HELLS-deficient cells displayed synergistic hypersensitivity in response to the combination treatment involving a non-toxic dose of MMS combined with clinically relevant dose of OLA in cell viability assays (**Fig. 3A**). To confirm these findings, we used live cell imaging (Incucyte) to monitor cell proliferation and survival over a period of 6 days. Cells were exposed to MMS+OLA for 1hr, followed by drug removal, and the addition of fresh growth media supplemented with OLA. We found that MMS+OLA was toxic to both HAP1 proficient and deficient cells inducing cell death in the first 2-3 days. However, HELLS proficient cells were more resistant and surviving cells recovered and proliferated reaching 60% confluence at 6 days, whereas HELLS deficient cells did not recover, and nearly all cells appeared to be effectively killed under these conditions (**Fig. 3C**). The significance of combining OLA with MMS, is highlighted by the fact that using both drugs together allowed for substantial reduction of the MMS dose from 100µM (**Fig. 3B**) to 2µM (50-fold decrease) (**Fig. 3A**) to achieve the effective killing of HELLS deficient cells.

To further investigate the relationship between PARP1 and HELLS, we used the PARP1 degrader molecule, iRucaparib-AP6 to degrade PARP1 in HAP1 cells and HAP1 HELLS KO cells. After 48hr, iRucaparib-AP6 induced efficient PARP1 degradation (**Fig. 3D**). The depletion of PARP1 in HAP1 cells resulted in a higher level of MMS sensitivity than the loss of HELLS alone (**Fig. 3D**). Notably, the combined loss of both HELLS and PARP1 further increased sensitivity to MMS suggesting that HELLS might compensate for the loss of PARP1 in mediating cell survival in response to DNA alkylation.

Since HELLS-deficient cells exhibit sensitivity to MMS and OLA treatment, we hypothesized that HELLS might promote the release of trapped PARP from chromatin. To test this hypothesis cells were subject to the standard PARP trapping treatment, involving exposure of cells to 1mM MMS combined with 10µM OLA for 1hr, followed by drugs removal and a recovery period of 15, 30, 60, and 120min (**Fig. S5A**). We found that PARP1 retention on chromatin increased 2-3 fold in both HELLS proficient and deficient cells after 1hr treatment, and PARP1 was progressively released from chromatin reaching nearly background levels by 120min post-treatment in HAP1 cells. HELLS-deficient cells appeared to have slightly elevated PARP1, especially during the early time points 15- and 30-min post-treatment (**Fig. S5B**). These data suggest that HELLS might contribute to the release of trapped PARP to some extent, however its loss does not appear to result in the major PARP trapping on chromatin.

In addition, we also tested the sensitivity of HeLa and CHON-002 HELLS deficient cells to OLA and a combination of OLA+MMS. We found that HELLS KO in HeLa and CHON-002 backgrounds were not sensitive to OLA or the combination of OLA and MMS (**Fig. S1C&D, F&G**). There are several reasons for the apparent lack of increased MMS+OLA sensitivity in HeLa and CHON-002 related to the abundance of endogenous HELLS and BER proteins. It was previously reported that MPG glycosylase levels play a significant role in mediating MMS sensitivity, where elevated MPG levels were driving increased sensitivity to MMS, whereas MPG downregulation was sufficient to substantially decrease MMS sensitivity [11, 56]. Indeed, the downregulation of MPG was sufficient to suppress MMS and Olaparib sensitivity of ALC1 deficient cells [68]. Therefore, it is reasonable to speculate that very low levels of MPG enzyme found in both HeLa and CHON-002 cells might partially contribute to the suppression of the MMS and Olaparib sensitivity phenotypes in HELLS KO cells. Furthermore, CHON-002 cells have substantially lower protein levels of HELLS and PARP1 as compared with HAP1 (**Fig. 2A**). In summary, these data suggest cell type-specific functions of HELLS in mediating the responses to DNA alkylation and imply the possibility of other pathways compensating for HELLS deficiency in mediating DNA damage repair.

### HELLS deficient cells accumulate elevated levels of Poly ADP ribose (PAR) upon DNA damage

It is well established that PARP1 is the key sensor of SSB and plays an important role in chromatin-based regulation of SSB repair through PAR-mediated chromatin reorganization and recruitment of DNA repair factors [18, 69]. In addition, rapid PARG1-mediated removal of PAR from chromatin is critical to ensure proficient and complete repair of SSB [70, 71]. To investigate the role of HELLS in SSB, we sought to determine if loss of HELLS impacts PARP1 activation in response to DNA alkylation by measuring PAR. To achieve this, cells were exposed to 500µM MMS for 30min with and without PARG inhibitor (PARGi) and OLA followed by the detection of PAR by immunofluorescence. We found that under normal growth conditions, HELLS KO cells have an elevated endogenous level of PAR compared to parental HAP1 cells. Elevated PAR could be the result of chronic activation of PARP1 by low levels of endogenous unrepaired SSB, consistent with deficient SSB. PAR polymer is unstable, therefore to enhance its detection, cells were treated with PARGi which inactivates the PARG1 enzyme and decreases its ability to degrade PAR. Indeed, we found a substantial increase of endogenous PAR in HELLS-deficient cells upon PAR stabilization with PARGi. Next, we determined the level of PAR in response to MMS-induced DNA damage and found a dramatic increase of PAR in the nuclei of the HELLS-deficient cells. To determine that the PAR increase was specifically due to enzymatic PARP activity, the cells were incubated with OLA, which substantially reduced the PAR signal, confirming PARP-mediated PAR formation and specificity (**Fig. 4A&B**). These data suggest that PARP is hyperactivated in response to MMS exposure in HELLS-deficient cells, which is consistent with high levels of unrepaired SSB inducing PARP activation.

**Figure 4.**
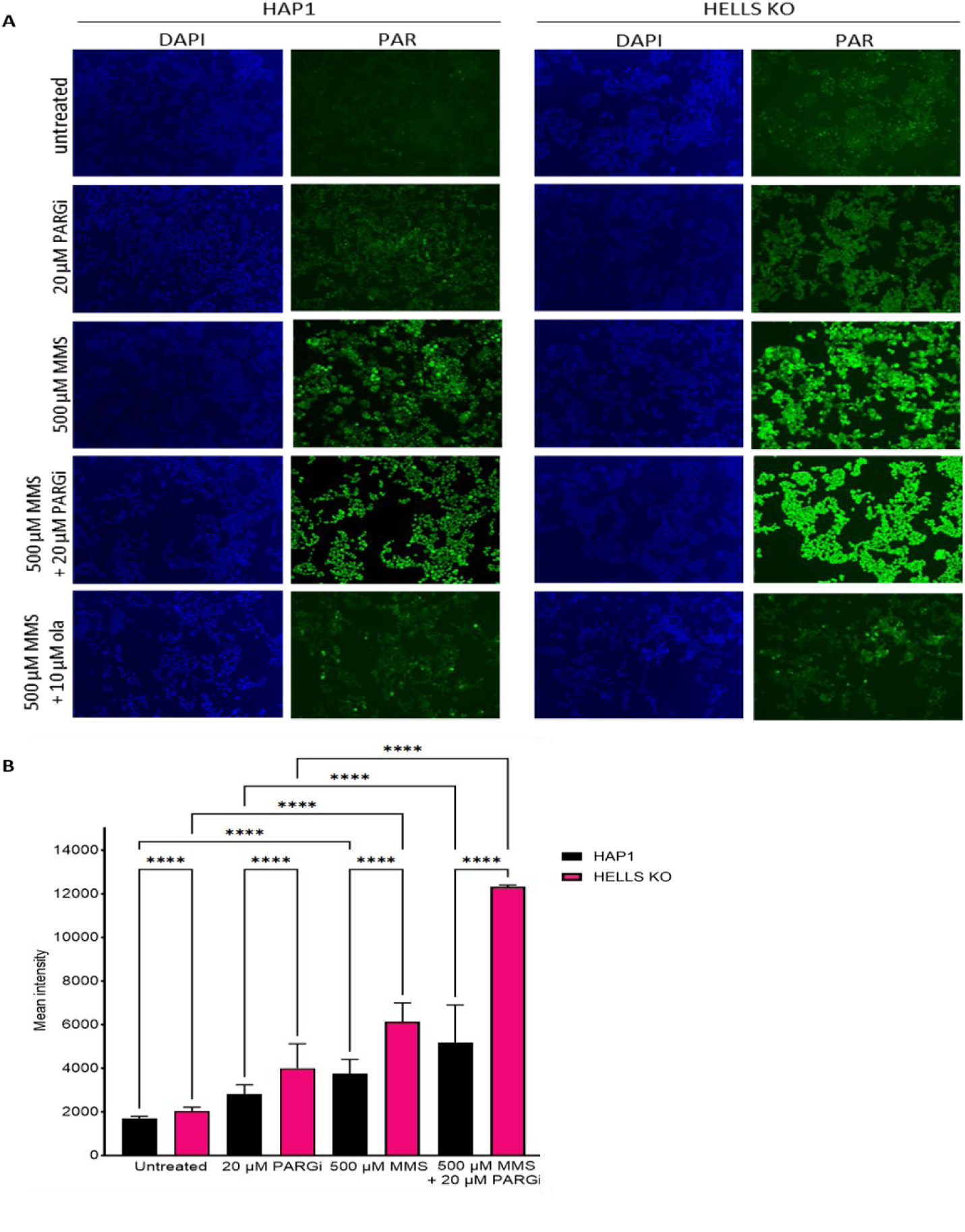
HELLS-deficient cells exhibit increased levels of PAR (Poly-ADP ribose). **A**. Representative images and **B**. fluorescence intensity of PAR upon indicated treatments in HAP1 and HAP1-HELLS KO cells. Data represent mean fluorescence intensity of 200 cells per sample of two independent biological experiments. Statistical significance was determined by two-way ANOVA with Tukey’s multiple comparisons test (****p <0.0001).

### HELLS and PARP1 are co-expressed in multiple cancer types

The significance of PARP1 in the DNA damage response (DDR), and as a key drug target in cancer is widely acknowledged [63, 69]. Since HELLS and PARP1 are known to be frequently overexpressed in many cancer types, we sought to determine if HELLS and PARP1 are co-expressed at the RNA and protein level by analyzing the TCGA and CPTAC web portals. We found that across all 33 human cancer types there is a positive correlation between HELLS and PARP1 mRNA expression with the highest co-expression correlation in GBM, TGCT, OV, LUSC, BRCA, and CHOL (**Fig. 5A**). Analysis of CPTAC revealed a positive co-expression correlation at the protein level in multiple cancer types including, GBM, LUAD, LSCC, COAD, and BRCA (**Fig. 5B**). The co-expression of HELLS and PARP1 in cancer implies potential functional interactions between HELLS and PARP1 in mediating chromatin remodeling and responses to DNA damage.

**Figure 5.**
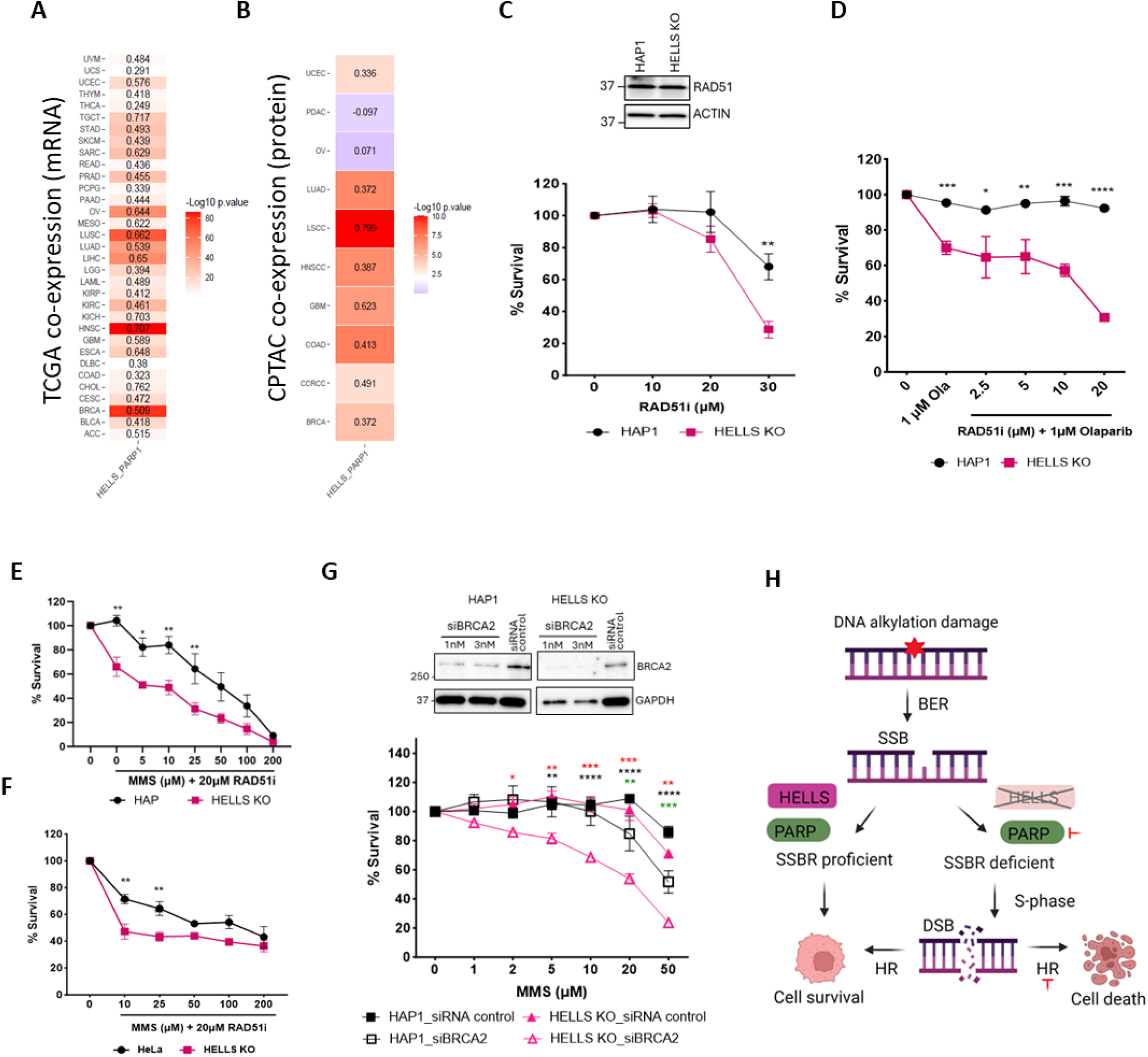
HELLS interacts genetically with PARP1 and HR pathway. **A**. HELLS and PARP1 co-expression was assessed based on RNA-seq data from TCGA. **B.** HELLS and PARP1 co-expression was analyzed based on proteomic data from CPTAC web portals. For each cancer type, the extracted, normalized expression levels for two genes (HELLS and PARP1) was determined. The co-expression and statistical significance were determined using the Pearson correlation coefficient analysis. **C.** Cell survival followed by RAD51 inhibition assessed by CCK8. HAP1 and HELLS KO cells were exposed to indicated doses of RAD51 inhibitor, B02, for 48hr. Western blot of WCE in HAP1 and HELLS KO cells probed for RAD51 indicating its basal expression level. **D**. HAP1 and HELLS KO cells were dosed with indicated concentrations of RAD51i and Olaparib for 48hr and cell survival assessed by CCK8. **E&F**. HAP and HeLa parental and HELLS KO cells were dosed with indicated doses of RAD51 inhibitor and MMS for 48hr and cell survival assessed by CCK8. All the cell survival data are mean ± SEM. n=3 independent biological replicates. **G.** Cell survival followed by BRCA2 downregulation assessed by CCK8. HAP1 and HELLS KO cells silenced with 1nM siRNA were dosed with indicated doses of MMS for 48hr. Asterisk colors denote the following comparisons: red-HAP1 siBRCA2 to HELLS KO siBRCA2; black-HELLS KO siRNA control to HELLS KO siBRCA2; green-HAP1 siRNA control to HAP1 siBRCA2. Western blot of WCE in HAP1 and HELLS KO cells probed for BRCA2 indicating downregulation of BRCA2 by siRNA for 6d. Statistical significance was determined by two-way ANOVA with Tukey’s multiple comparisons test (*p<0.05, **p<0.01, ***p<0.001, ****p<0.0001). **H.** Proposed model of HELLS mediated repair of SSBs in response to DNA alkylation damage.

### Loss of HELLS is synthetic lethal with HR inhibition

The HR pathway modulates cell survival in response to DNA alkylation damage [11]. Furthermore, it is well established that PARP inhibition is synthetic lethal with HR deficiency [63]. Previous studies in fungal cells reported synthetic growth defects in fungal cells where loss of MUS30 (HELLS) was combined with loss of MEI3 (RAD51) inducing HR pathway deficiency [40]. We sought to determine if loss of HELLS is synthetic lethal with HR inhibition in human cells. We used the B02, a potent RAD51 inhibitor that disrupts RAD51 binding to DNA, and blocks HR-mediated repair of DSB during S-phase [72, 73]. We found that HAP1 HELLS KO cells display increased, dose-dependent sensitivity to the RAD51 inhibitor (**Fig. 5C**). This sensitively was further potentiated when RAD51i was combined with OLA, or MMS (**Fig. 5D&E**). These data suggest that loss of HELLS potentiates synthetic lethality between PARP inhibition and HR deficiency. In addition, we found that HELLS-deficient HeLa cells displayed enhanced sensitivity to RAD51i upon DNA alkylation damage (**Fig. 5F**). Consistently, we also found that BRCA2 downregulation potentiates the sensitivity of HELLS KO cells to DNA alkylation damage, providing additional evidence that RAD51 activity mediates cell survival in HELLS-deficient cells (**Fig. 5G**). Taken together, these data suggest that the HELLS protein functions outside of the canonical HR pathway, and the activity of the RAD51/BRCA2-mediated HR pathway compensates for the loss of HELLS in support of cell survival in response to DNA alkylation damage. We propose a model where HELLS functions as an important regulator of SSB repair and mediates cell survival in response to DNA alkylation damage. In cells that are HELLS and PARP deficient, SSBR is compromised, and therefore these cells must heavily rely on proficient HR for survival (**Fig. 5H**).

### Loss of HELLS does not affect expression of most DDR genes, but impacts cell differentiation genes

To examine the effect of HELLS loss on the expression of DNA Damage Response (DDR) and related pathways, we performed RNA-seq in HAP1 and HELLS KO cells, as well as analyzed publicly available RNA-sequencing data for HeLa and HeLa HELLS KO cells (GEO, GSE 136931). We observed that only 8 (FMN2, CDA, PDE4B, AK1, TWIST1, APLF, GGN, EID3) out of 464 DDR genes showed a significant change in expression at an FDR-corrected p-value cut-off of 0.05 and log 2-fold-change cut-off of 1, suggesting that HELLS KO does not substantially influence the expression of DDR genes in the HAP1 or HeLa cell lines **(Fig. 6A**). We performed a gene ontology enrichment analysis to determine the biological processes affected by HELLS KO in HAP1 cells. We observed that the upregulated genes (N=295) were mainly associated with RNA metabolism, and the downregulated genes (N=309) were related to cell development and differentiation **(Fig. 6B**). To further interpret the transcriptional effects of HELLS KO, we performed Transcription Factor (TF) enrichment analysis (TFEA) for the upregulated and downregulated genes. This analysis prioritizes TF based on the overlap between given lists of differentially expressed genes and previously annotated TF targets assembled from multiple publicly available resources, including ChIP-seq experiments from ENCODE, ReMap, and individual publications. Many top-ranked (or enriched) TFs were also differentially expressed in the HELLS KO samples. For example, PRRX1 was identified as one of the top TFs associated with downregulated genes and exhibited significant downregulation of expression in HAP1 HELLS KO cells (**Fig. 6C**). This is a notable finding since PRRX1 is known to facilitate epithelial-to-mesenchymal transition (EMT), and metastasis of many cancers [74, 75]. This data suggests that targeting HELLS could potentially slow down cancer progression in certain contexts by reducing the expression of transcription factors associated with the differentiation state of tumor cells and EMT.

**Figure 6.**
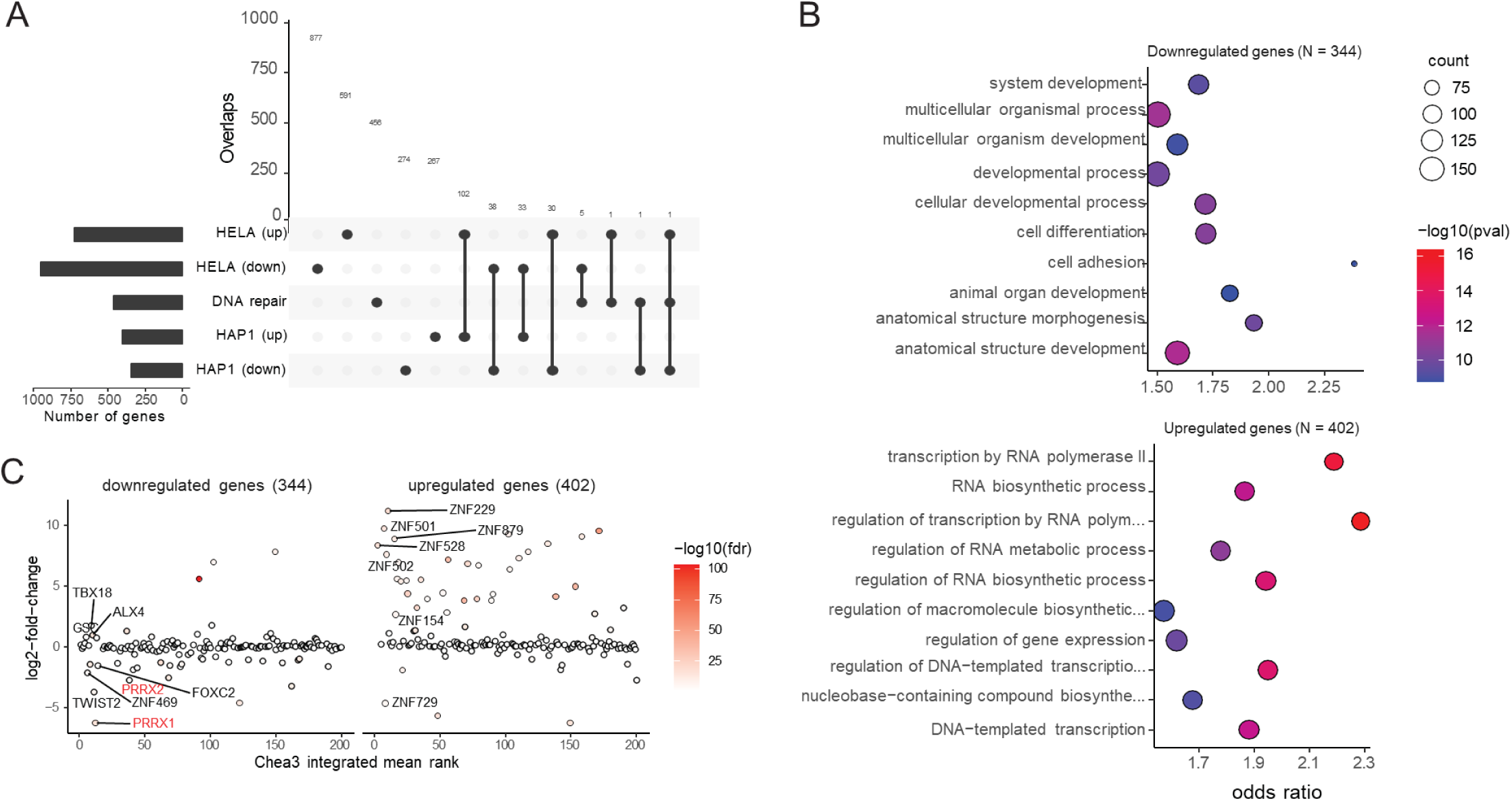
Transcriptional profile of HELLS KO cells. **A.** Upset plot showing the overlap between upregulated and downregulated genes from HELLS KO in HAP1 and HeLa cells and 464 genes in DNA Damage Response and related pathways (i.e., Hallmark DNA repair, Double-strand break repair, and Base excision repair pathways), **B.** Gene ontology analysis showing enrichment of biological processes associated with upregulated and downregulated genes in HAP1 cells. **C.** Scatter plot showing the association between log2 fold-change in expression and integrated mean rank for TFs for upregulated and downregulated genes in HAP1 cells.

## Discussion

The potential for SNF2 chromatin regulators to modulate the sensitivity of tumors to existing anticancer drugs, including PARP inhibitors presents new avenues of clinical investigation [4, 68, 76, 77]. The HELLS SNF2 family remodeler has been implicated in the regulation of key cancer-driving pathways, including proliferative signaling, oncogenic transcriptional programs, and DNA damage response [4, 7]. Our studies uncover a new regulatory link between HELLS and the repair of SSB in mediating responses to clinically relevant DNA alkylation damage in cancer cells. We found that loss of HELLS in HAP1 cells results in deficient SSB repair and selective sensitivity of cells to DNA alkylating agents and PARP inhibitors. Furthermore, HELLS-deficient cells display synthetic lethality with inhibition of RAD51, highlighting the potential therapeutic value of targeting HELLS in cancer.

The SSBs are among the most abundant DNA lesions in mammalian cells, arising in the order of tens to hundreds of thousands per cell per day [78]. The levels of SSBs in cancer cells are likely much higher due to metabolic dysregulation and substantial increase in ROS-induced DNA damage [79]. Unrepaired SSBs have the potential to generate high levels of replication-associated DSBs, arising when replication forks collide with unrepaired SSBs, resulting in replisome disassembly and fork collapse [21].

The chromatin-based regulation of SSB repair is not well understood, and BER/SSBR is known to be frequently dysregulated in cancer [80, 81]. Our studies demonstrate that loss of HELLS in cancer cells phenocopies several key characteristics associated with SSB deficiency. We found that HELLS-deficient cells display decreased cell proliferation, increased sensitivity to DNA alkylating agents, and PARP inhibitors, G2/M cell cycle arrest, elevated PAR, increased formation of micronuclei in response to DNA alkylation, and dependence on proficient HR for cell survival. We found that loss of HELLS substantially impairs enrichment of the key BER proteins mediating SSBR, including PARP2, XRCC1, POLB and LIG3 (**Fig. 2I**) on damaged chromatin, suggesting HELLS plays a role in coordinating the recruitment and/or retention of DNA repair proteins in response to MMS-induced DNA damage. Previous studies have reported a role for HELLS in loading macroH2A histone variant onto chromatin, and macroH2A has been implicated in promoting BER [82–84]. Although we have not detected changes in global levels of chromatin-associated macroH2A in HELLS KO cells, it is possible that HELLS may promote some aspects of BER via its role in macroH2A deposition at specific genomic domains. Collectively, our studies reveal a new role for HELLS in BER/SSBR pathway, and align with previous studies reporting the association between HELLS, macroH2A and BER.

Notably, we determined that ATP-dependent activity of HELLS is necessary to promote SSBR, which is consistent with its chromatin remodeling function (**Fig. 2H**). At the organismal level, single-strand break (SSB) deficiency is linked to various conditions including developmental defects, neurological disorders, and premature aging [1]. Consistently, mice deficient in HELLS (LSH) exhibit severe developmental growth defects and premature aging phenotypes associated with replicative senescence [85]. Our studies demonstrate the selective sensitivity of HAP1 HELLS-deficient cells to clinically relevant drugs that induce the formation of SSB, including DNA alkylators and PARP inhibitors (**Fig. 1A-F**, **Fig. 3A-C**). Importantly, our data imply cell type specificity of HELLS function suggesting that highly proliferating cancer cells with elevated HELLS expression might rely on HELLS for SSB repair, whereas HELLS might be dispensable in slowly proliferating, non-transformed cells with low HELLS expression (**Fig. 1A-F**, **Fig. 2A, Fig. S1A, E,F,G**). Our data suggests that ATPase activity of HELLS is required for SSBR, but chromatin remodeling appears to be dispensable for cell survival in response to MMS in HAP1 cells, since HELLS K254R (ATP-ase inactive) mutant cells do not display increased sensitivity to MMS (**Fig.S4A**). The lack of MMS sensitivity, despite deficient SSBR could be explained by the compensatory mechanisms operating at the replication fork involving robust RAD51-mediated fork protection, and HR of SSB-derived DSB. Indeed, we observed that HELLS K254R mutant cells lack MMS sensitivity, but display increased sensitivity to combination treatment when MMS is combined with RAD51i (**Fig.S4B**). Similarly, we have observed that HELLS KO HeLa cells demonstrate deficient SSBR, but lack obvious sensitivity to MMS alone (**Fig.S1B**). However, inhibition of RAD51 activity renders HeLa HELLS KO cells sensitive to MMS (**Fig.5G**). Alternatively, it is reasonable to suggest that HELLS could have dual role, involving chromatin remodeling-dependent functions during SSBR, and chromatin-remodeling-independent functions, operating during the replication fork protection.

Indeed, recent study showed that HELLS maintains fork protection via macroH2A deposition and RAD51 filament formation at stalled replication forks, revealing that HELLS-deficient U2OS cells displayed impaired RAD51 deposition at stalled replication forks [38]. Consistently, in our studies we have found that RAD51 accumulation on chromatin in response to MMS was decreased in HELLS KO cells (**Fig. 2I**), pointing to an important co-operation between HELLS and RAD51 in response to MMS-induced replication stress. Interestingly, it was previously reported that the catalytically inactive K254R HELLS mutant protein was able to associate with chromatin and nucleosomes [36, 84] and it was also able to physically associate with macroH2A histone variant in vitro [84]. Therefore, it is possible that the association of K254R HELLS with nucleosomes, and macroH2A is sufficient for RAD51-mediated fork protection and cell survival in response to MMS-induced replication stress.

Prior animal studies demonstrated that sensitivity to MMS-induced DNA alkylation varies greatly across different organs, tissues, and cell types, and was shown to be significantly modulated by the levels and activity of MPG (AAG) enzyme that initiates BER of alkylation DNA damage [11]. In addition, recent studies reported that MPG enzyme was the key mediator of MMS, PARPi sensitivity, and synthetic lethality with HDR in ALC-/- cells [68, 77]. These observations raise the possibility that MPG levels might also determine sensitivity phenotypes in HELLS-deficient cells. Indeed, we found that MPG levels are higher in HAP1 cells, as compared to HeLa and CHON-002 (**Fig. 2A**). Therefore, it is tempting to speculate that low MPG might partially suppress the initial formation of SSB, and as a result suppress MMS and PARPi sensitivity phenotypes in HeLa and CHON-002 HELLS deficient cells (**Fig. S1B-G**). Furthermore, the functional status of HR has been shown to determine the survival outcomes in response to DNA alkylation damage and PARP inhibitors [11, 56, 59].

Our studies imply that HELLS does not appear to directly modulate the repair of alkylation-derived DSB in HAP1 cells, since both HELLS proficient and deficient cells exhibited equal levels and progressive formation of alkylation-derived DSB during 0-6hr post-MMS exposure (**Fig. 2D**). Significantly elevated DSB at 24rh post-MMS in HELLS-deficient cells likely represent DSB that arise during S-phase, as a result of replication fork collapse upon collision with unrepaired SSBs (**Fig. 2B-D**), and may also include spontaneously generated DSB as part of apoptosis process, rather than they are the indication of deficient HR. This is further supported by the lack of substantial sensitivity in HELLS-deficient cells to the DSB-inducing agents, etoposide and topotecan (**Fig. 1F&G**). These results are consistent with several prior studies reporting that the overall capacity of canonical HR was not affected in HELLS-deficient U2OS, MCF10A and HEK293T cell backgrounds [36, 38, 57]. However, it should be pointed out that HELLS was found to facilitate repair of IR-induced two-ended DSB repair within heterochromatic regions during G2 phase of the cell cycle through interactions with CtIP [36]. Collectively, multiple independent studies highlight the context and cell type-specific functions of HELLS in DNA repair.

Our findings that HELLS-deficient HAP1 cells exhibit increased sensitivity to PARP inhibition (**Fig. 3A-C**) are consistent with several CRISPR screen-based studies identifying HELLS as a mediator of PARPi sensitivity in breast, ovarian, and prostate cancer [64–66]. Although HR deficiency is known to confer sensitization to PARP inhibitors, several studies have shown that SSBR deficiency can substantially sensitize cells to PARP inhibitors [68, 86, 87]. Indeed, cells deficient in the key SSBR proteins such as POLβ, and XRCC1 display hypersensitivity to PARP inhibitors [14]. DNA alkylators, including clinically used Temozolomide, can potentiate cytotoxicity of PARPi [88–90]. Our findings that HAP1 HELLS-deficient cells are hypersensitive to DNA alkylators in combination with PARPi (**Fig. 3A-C**) are consistent with other studies where increased SSBs and PARP trapping on chromatin cause generation of S-phase-dependent DNA damage leading to cell death [23, 91].

Our findings that loss of HELLS is synthetically lethal with HR inhibition are consistent with the well-established synthetic lethality relationship between PARP inhibition and HR deficiency, where unrepaired SSB are converted to lethal DSB during S-phase in the absence of functional HR. The HELLS-deficient cells, when subjected to HR inhibition, exhibit a similar vulnerability, providing additional evidence supporting the role of HELLS in SSB repair. The observed synthetic lethality with HRD, and selective sensitivity of HAP1 HELLS deficient cells to DNA alkylators and PARP inhibitors, but not DSB-inducing agents, phenocopies loss of ALC1, another SNF2 remodeler implicated in promoting BER and release of trapped PARP [68, 77, 86]. These studies underscore the important role of SNF2 chromatin remodeling enzymes in promoting efficient BER/SSBR in the context of chromatin.

Considering the substantial overexpression of HELLS in various tumor types, it is possible that HELLS in addition to promoting cancer cell proliferation, may also aid tolerance of elevated ROS-induced DNA damage and replication stress. We found that overexpression of HELLS in HeLa HELLS KO resulted in robust SSBR and increased resistance to MMS (**Fig. 2G**). The co-expression of HELLS and PARP1 in multiple cancer types (**Fig. 5A**) is significant and implies that HELLS might cooperate with PARP1 in regulating various aspects of DNA damage response and/or DNA damage tolerance, promoting cancer cell proliferation and survival despite the elevated DNA damage.

Alkylating agents and PARP inhibitors are widely used in cancer therapy [63, 88, 90]. Therefore, targeting HELLS in cancer could potentially lead to a synergistic response of halting rapid cell proliferation and hyper-sensitizing cancer cells to chemotherapy, and/or PARP inhibitors. In addition, it would be important to assess if HELLS deficient cancer types are hypersensitized by a combination of PARPi and DNA alkylation, or RAD51 inhibitors. Furthermore, HELLS targeting could potentially offer a specific vulnerability in tumors with HR deficiency.

The PRRX1 and PRRX2 transcription factors, play significant roles in cancer by promoting epithelial-mesenchymal transition (EMT), a process crucial for cancer metastasis [74, 92–94]. Elevated levels of PRRX have been associated with increased tumor invasiveness and poor prognosis in various cancers [74, 75, 92–96]. Our study identifies HELLS as an upstream regulator of PRRX1 and PRRX2 in leukemia HAP1 cells (**Fig. 6C**), raising the possibility that targeting HELLS could potentially reduce EMT and tumor invasiveness through the downregulation of PRRX in certain contexts.

In conclusion, our study underscores the important role of HELLS in the repair of SSBs and mediating responses to DNA alkylation damage in certain cellular contexts. The synthetic lethality between HELLS deficiency and HR inhibition, combined with the heightened sensitivity to PARPi and DNA alkylating agents, provides new insights into chromatin-based regulation of DNA repair in cancer, which could lead to new avenues for targeted and more effective cancer therapies.

## Acknowledgments

We thank members of the Czaja lab for discussions and suggestions and Hiruni Marsha Aponso for technical assistance. This work was supported by NIH grant R01GM143428 (to W.C). A.D. was supported by NIH grant R01CA060499 (to A.D.). Z.A.L. was supported by NIH grant R01GM143428 (to W.C). S.A.G. was supported by NIH grant R01DK132781 (to S.A.G.). R.C-G. was supported by NIH grant R01GM143428 (to W.C) and NIH/NCI grant R21CA259630 (to R.C-G), and the National Scleroderma Foundation (to R.C-G.). We thank the UAB Flow Cytometry and Single Cell Core Facility for the use of the facility.

The content is solely the responsibility of the authors and does not necessarily represent the official views of the National Institutes of Health.

## Author contributions

W.C. designed the study with contributions from J.T.J., C.M.W. and EP. W.C., S.A.G., Z.A.L., and A.D. provided resources. J.T.J., C.M.W., E.P., A.K., Y.L., J.S.R.N., O.D., K.R.K., A.I., E.S., R.C-G, A.D., and W.C. designed and conducted experiments. B.A.J. and A.S. performed the RNA-seq analysis. Y.Z. performed analyses of TCGA and CPTAC databases. W.C. wrote the manuscript with contributions from J.T.J., C.M.W., and E.P. All authors had editorial input.

## Resource Availability

Lead Contact: Wioletta Czaja

Requests for further information and resources should be directed to and will be fulfilled by the lead contact, Wioletta Czaja.

## Materials availability

All unique/stable reagents generated in this study are available from the lead contact without restriction.

## Data and code availability

All data and code reported in this paper will be shared by the lead contact upon request.

## Declaration of generative AI and AI-assisted technologies in the writing process

During the preparation of this work the author(s) used Grammarly and ChatGPT in order to improve language and readability. After using this tool/service, the author(s) reviewed and edited the content as needed and take(s) full responsibility for the content of the publication.

## MATERIALS AND METHODS

### DNA damaging drugs and chemicals

Methyl methanesulfonate (MMS), temozolomide (TMZ), and the Rad51 inhibitor B02 were obtained from Millipore Sigma. Etoposide was purchased from Cayman Chemical. Olaparib was obtained from Fisher Scientific. The PARG inhibitor PDD00017273 and iRucaparib-AP6 were obtained from MedChem Express.

### Cell lines and Cell Culture

The human near-haploid chronic myelogenous leukemia-derived HAP1 cell line (C631) and HAP1 HELLS knockout (HZGHC001202c003) containing a single base pair insertion in exon 3 were purchased from Horizon Discovery and maintained in IMDM (Gibco) media containing 10% Fetal Bovine Serum (Gibco) and 1x penicillin/ streptomycin (Corning). Parental HeLa, and HeLa HELLS knockout cells (ab65006, Abcam) containing a 1 base pair insertion and a selection cassette in exon 2 were obtained from Abcam and were cultured in DMEM (Gibco) media containing 10% Fetal Bovine Serum and 1x penicillin/ streptomycin. U2OS (ATCC), and MCF7 (ATCC) cells were cultured in DMEM media supplemented with 10% Fetal Bovine Serum and 1x penicillin/ streptomycin. CHON-002 cells were cultured in DMEM media supplemented with 10% heat-inactivated Fetal Bovine Serum and 1x penicillin/ streptomycin. The HELLS knockout in CHON-002 cells, containing a single base pair insertion in exon 3, was created using vector plentiCRISPR V2 containing a guide RNA targeting the sequence ATAGAGAGTCGACAGAAATT in exon 3. All cell lines were cultured at 37°C and 5% CO2. Routine mycoplasma testing was performed using the Southern Biotech mycoplasma detection kit.

### Western blot

HAP1 and HELLS KO cells were lysed in RIPA buffer (ChemCruz) containing HALT protease inhibitors (Thermo Scientific), phosphatase inhibitors A and B (Santa Cruz), and PMSF for 30 min on ice. Total protein concentration was determined using BCA (Thermo Scientific). The lysates were resolved on 4-15% Acrylamide TGX gel (Bio-Rad) by SDS-PAGE and the proteins were transferred to LF-PVDF membrane (Bio-Rad). The membrane was blocked with Prometheus OneBlock (Genesee Scientific) and probed with primary antibodies overnight at 4°C. Western blots utilized the indicated antibodies: HELLS (Cell Signaling), Actin (SigmaAldrich), GAPDH (Cell Signaling), PARP1 (Cell Signaling), PARP2 (Santa Cruz Biotechnology), PARG (Cell signaling), γ-H2AX (Cell Signaling), Rad51 (EMD Millipore and Abclonal), pATM (Santa Cruz), MPG (Abcam), APEX (Abclonal Technology), XRCC1 (Santa Cruz Biotechnology and Active Motif), XRCC5 (Abclonal Technology), DNA LIG III (Santa Cruz Biotechnology), POLβ (Abcam), MacroH2A (Abcam), and PAR binding reagent (MilliporeSigma). The following day, the membranes were incubated with HRP-conjugated secondary antibodies for 2 hr, washed and visualized using Clarity ECL (Bio-Rad) with a ChemiDoc MP (Bio-Rad).

### Apoptosis Assay

A total of 250,000 HAP1 or HELLS KO cells were seeded in 6-well plates. After two days of growth, cells were treated with the indicated concentration of MMS for 1 hour. Following treatment, the cells were washed with 1x PBS, fresh media was added, and the cells were incubated overnight. The following day, all cells were harvested and stained with propidium iodide and FITC Annexin V (BD Biosciences) according to the manufacturer’s protocol. Apoptotic and necrotic cell death was assessed using an LSR Fortessa (BD Biosciences), measuring Annexin V and propidium iodide signal intensities. Total apoptotic death was determined by combining early apoptotic (quadrant 4) and late apoptotic/necrotic (quadrant 2) populations.

### Global DNA methylation ELISA

For Global 5-mC DNA methylation (5-mc) ELISA, genomic DNA was prepared from the parental and HELLS KO cells of HAP1 and HeLa using PureLink genomic DNA prep kit (Invitrogen). Global 5-mC DNA methylation levels were quantified using the 5-mC DNA Methylation Colorimetric Assay Kit (Abcam) according to the manufacturer’s instructions using 100ng of input DNA with 260/280 ratio >1.6.

### Comet Assay

Comet assays were performed following the MIRCA guidelines [97]. For the alkaline comet assay, 400,000 parental HAP1 cells and HELLS KO cells were seeded in 6-well plates and grown overnight. Cells were treated with 500 µM MMS for 1 hr, then harvested at 0, 3, 6, and 24 hr post-treatment. After harvesting, cell pellets were washed with ice-cold PBS and resuspended at a concentration of 1x10^6^ cells/mL in ice-cold PBS. After harvest, cells were either immediately plated on slides precoated with 1% agarose or stored at 4°C for up to 6 hr while other timepoints were collected before plating. All reagents were from the OxiSelect Comet Assay Kit (Cell Biolabs). Briefly, 10µL cells were resuspended in 90µL fresh low melt agarose. The slides were immersed in lysis buffer and were incubated overnight at 4°C. Before electrophoresis, the samples were incubated in pre-chilled alkaline solution for 30 min. Electrophoresis was conducted at 1 V/cm and 300 mA for 40 min in chilled electrophoresis buffer. For other cell lines, the assay was similarly performed with minor modifications. Specifically, 200,000 parental HeLa and HELLS KO cells were treated with 1 mM MMS and harvested the pellets were resuspended at 5x10^5^ cells/mL. In CHON-002 cells, 200,000 parental and HELLS KO cells were grown for three days before treatment with 500 µM MMS. For experiments with MCF7 cells, 150,000 cells were reverse transfected with the indicated siRNA. After 72hr, cells were treated with 1 mM MMS before harvesting. Alternatively, 100,000 U2OS cells were reverse transfected with the indicated siRNA. After 48hr, cells were treated with 1 mM MMS. All cells were harvested at the indicated timepoints. Electrophoresis was performed for 20 min in HeLa cells and 30 min in CHON-002, MCF7, and U2OS cells. All slides were stained with Vista Green DNA Dye for 1 hr, and images were captured using an EVOS Fluorescence Microscope (AMG), with subsequent analysis performed using CometScore software. For the neutral comet assay the cells were grown, treated and prepared on slides as in the alkaline method. The slides were then submerged in a neutral lysis solution (2% SDS, 0.5 M Na2EDTA, 0.5 mg/mL Proteinase K, pH 8.0) at 37°C overnight. Slides were transferred to a neutral rinse/electrophoresis solution (90 mM Tris buffer, 90 mM boric acid, 2 mM Na2EDTA, pH 8.5) for 30 min before being placed in an electrophoretic chamber with the neutral buffer. Electrophoresis was performed at 1 V/cm and 300 mA for 30 min at room temperature. The slides were visualized and quantified as described above.

### Cell Transfection and RNA Interference

HeLa HELLS KO cells (400,000 cells per well) were seeded in 6-well dishes and incubated overnight. Cells were transfected with a pCMV-HELLS (Origene). Transfection was performed using Lipofectamine 3000 (Invitrogen) according to the manufacturer’s instructions. For RNA interference 5,000 cells per well (96 well plate) or 80,000 cells per well (6 well plate) of HAP1 and HELLS KO cells were transfected with 1nM or 3nM silencer select BRCA2 siRNA (Ambion). Alternatively, 150,000 MCF7 cells per well or 100,000 U2OS cells per well (6 well plates) were transfected with 3nM silencer select HELLS or negative control siRNA (Ambion). Reverse transfection was performed using RNAiMAX (Invitrogen) according to the manufacturer’s instructions.

### Cell Cycle Analysis

Asynchronous cells were treated with 0, 250 or 500 µM MMS for 1hr and allowed to recover for the indicated time. Cells were fixed with 70% EtOH, and treated with propidium iodide and RNAse A. Cell cycle distribution was determined by flow cytometry using an LSR Fortessa (BD Bioscience). Quantification was performed in FlowJo. All experiments were performed in at least triplicate.

### Live-cell imaging assay

HAP1 parental and HELLS KO cells (5,000 cells per well) were seeded in 96-well plates continuously monitored utilizing the IncuCyte live-cell analysis system (Essen BioScience, Ann Arbor, MI, USA). Cells were maintained at 37°C and 5% CO2. Data was processed and analyzed using IncuCyte S3 software.

### Cell Survival Assays

CCK-8 cell survival assays were performed by seeding 15,000 HAP1 cells, 4,000 HeLa cells, or 10,000 CHON-002 cells in 96 well plates and adding drugs after 24 hr. Viability was assessed 2 days after treatment. In proliferation assays 5,000 HAP1 cells or 2,000 HeLa cells were plated, and growth was assessed at 16, 24, 48, 72, and 96 hr. CCK-8 assay (Dojindo) was performed following the manufacturer’s guidelines. For crystal violet-based cell survival, 100,000-150,000 HAP1 or HELLS KO cells were plated in 6 well dishes. The following day, the cells were treated with the indicated concentration of MMS, TMZ, Olaparib, or Etoposide. After a 3-day incubation with the compounds, the cells were washed, fixed with 10% formalin for 30 min, and stained with 0.01% crystal violet for 30 min. The excess crystal violet was removed with 2 washes of 1x PBS and the plates were imaged on a ChemiDoc. For quantification, the crystal violet was solubilized with methanol. 50-100uL of the solubilized crystal violet was plated in duplicate in a 96-well plate and absorbance was read at 570nm. All experiments were performed in at least triplicate. For colony formation assays 300 cells were seeded in 6 well dishes. The following day, the cells were dosed with the indicated concentration of MMS or Olaparib. After 10 days, the colonies were stained as above.

### Chromatin Fractionation and PARP trapping

HAP1 and HELLS KO cells were plated in 6 well dishes. After 48 hr, the cells were treated with 1mM or 3mM MMS for the indicated time. PARP trapping experiments also included 10 µM Olaparib. The media was removed, and the cells were washed with 1xPBS. Fresh media was added to the cells, and they were allowed to recover for up to 2hr. Multiple wells were harvested for each condition at the indicated timepoints. The cells were lysed and fractionated using the subcellular fractionation kit (Thermo Scientific) according to the manufacturer’s directions. 10 µg of each fraction was resolved by SDS-PAGE and Western blots were performed as described above.

### RNA-seq and data analysis

Duplicates of RNA samples from HAP1 and HELLS KO were prepared for RNA-seq. Total RNA was prepared from 1x10^6^ cells of HAP1 and HELLS KO using Direct-zol RNA Microprep kit (Zymo Research). The RNA was checked for quality using the Agilent BioAnalzyer 2100. The libraries were made with the NEB Next Ultra II Directional RNA Seq kit following the manufacturer’s protocols. The resulting libraries were sequenced on the NovaSeq 6000 with Paired-end 100bp chemistry. Sequencing produced approximately 40M reads per sample. RNA-seq data from HeLa was downloaded from GEO (GSE 136931). The raw data was aligned to the GRCh38 version of the human genome using the STAR (v2.7.3a), and non-duplicated reads with mapping quality ≥ 20 were retained for downstream analysis. RNA-seq counts for genes were generated using the Subread featureCounts function (v2.0.6). Differential expression analysis was performed using DESeq2(v1.36). Gene ontology analysis was performed using topGO (v2.48), and transcription factor enrichment analysis was performed using ChIP-X Enrichment Analysis Version 3 or ChEA3.

### Cell sorting

Single-cell sorted diploid clones were obtained using a FACSymphony S6 Sorter (BD). 10 million parental and HELLS KO HAP1 cells were stained with Hoechst (Life Technologies) for 1 hr. Cells exhibiting the 4N peak were sorted into single wells of a 96-well plate and amplified. To generate revertants, HELLS-KO HAP1 cells were passaged at least 10 times after receipt from Horizon.

### γH2AX Immunofluorescence

Cells were seeded and grown on four-well chamber slides for 24 hr. Following that time, the cells were incubated for 1 hr in 250 or 500 µM MMS and the immunofluorescence protocol was performed at different times post MMS treatment. Cells were washed with PBS twice, fixed with ice-cold methanol for 5 min, and washed with PBS Tween 0.1% (PBST) three times. After blocking with 3% bovine serum albumin in PBS for 1 hr at room temperature, cells were incubated with anti-rabbit γH2AX (1:400; Cell Signaling), overnight at 4°C. Slides were washed with PBST three times and incubated with Alexa Fluor 594 conjugated goat and/or Alexa Fluor 555 conjugated goat secondary antibodies (1:200 Invitrogen) for 1 hr. The nuclei were stained with DAPI and the slides were visualized and documented using an IX73 inverted fluorescent microscope connected to an XXD DP80 camera. The number of dots per cell were quantified using Cell Profiler Software. Fluorescent intensity was analyzed using the Cellsens Dimension Desktop Version 4.2 software. Statistical analysis was performed using one-way ANOVA with multiple comparisons.

### PAR Immunofluorescence

200,000 HAP1 and HELLS KO cells were seeded on four-well chamber slides. After 24 hr, cells were exposed to 500 µM MMS, 20 µM PARGi, 500 µM MMS + 20 µM PARGi, or 500 µM MMS + 10 uM Olaparib for 30 minutes. Cells were washed twice with PBS and fixed with 4% paraformaldehyde for 20 minutes. The samples were blocked with immunofluorescence blocking buffer (Cell Signaling) before overnight incubation in anti-mouse PAR primary antibody (1:100; Santa Cruz) in antibody dilution buffer (1% BSA and 0.1% Tween 20 in PBS). PAR was detected with Alexa Fluor 488 conjugated goat anti-mouse secondary antibody (1:150; Abcam) and the nuclei were stained with DAPI. The slides were visualized and documented using an IX73 inverted fluorescent microscope connected to a XXD DP80 camera.

### Micronuclei assay

Cells were exposed to indicated doses of MMS for 1 hr followed by drug removal and cell recovery in the incubator for 48 hr. Cells were treated with 10 µg/mL KaryoMAX™ Colcemid™ Solution (Thermo Fisher Scientific) for 16 hr. Cells were trypsinized, suspended in 0.56% KCl hypotonic solution for 15 min at 37 °C, and then resuspended in hypotonic solution with 0.05% Tween. To make chromosome spreads and detect chromosomes and micronuclei, cells were spotted onto slides by centrifugation at 290 × g for 5 min using a cytospin centrifuge. IF was done by fixing the chromosome spreads with 4% paraformaldehyde (PFA) for 10 min. The chromosome spreads were then counterstained with DAPI solution ProLong® Gold Antifade Mountant with DAPI (Thermo Fischer Scientific) and visualized in a Zeiss LSM 900 live cell confocal microscope with Airyscan 2 system. Quantitation of fluorescent images was performed with ImageJ. The statistical significance of micronuclei and ploidy numbers was calculated using one-way ANOVA with multiple comparisons.

### Plasmid cloning and Stable cell line construction

Untagged HELLS in the pCMV6-XL4 vector (SC126963) and Myc-DDK HELLS in the pCMV6-Entry vector (RC212231) were purchased from Origene. A custom mutation of the Myc-DDK HELLS construct that encodes HELLS K254R was created by Origene. pLenti V HELLS WT or Mutant plasmids were constructed using In-Fusion cloning (Takara Bio) according to the manufacturer’s instructions. HELLS cDNA was amplified by PCR using Q5 High-Fidelity DNA Polymerase (NEB). Primers used to amplify HELLS cDNA were F (5’TCGTGACGCGGGATCCGCCACCATGCCAGCGGAACGGCCC) and R (5’CGCCGCTGCCGCTAGCAAACAAACATTCAGGACTGGAATCTTCAG). LentiV_Cas9_puro (Addgene) was digested with BamH1 and Nhe1. To package lentivirus, pLenti V HELLS was co-transfected with psPAX2 and pMD2G plasmids into HEK293T cells using 3 µg/mL Polyethylenimine Max (Polyscience). 1 mL of virus supernatant was transduced with 8 µg/mL of polybrene (Millipore Sigma) onto 1x10^6^ HAP1 cells in a 6 cm dish overnight. Cells were selected with 1 mg/mL Puromycin (Millipore Sigma).

## SUPPLEMENTARY DATA

**Figure S1.**
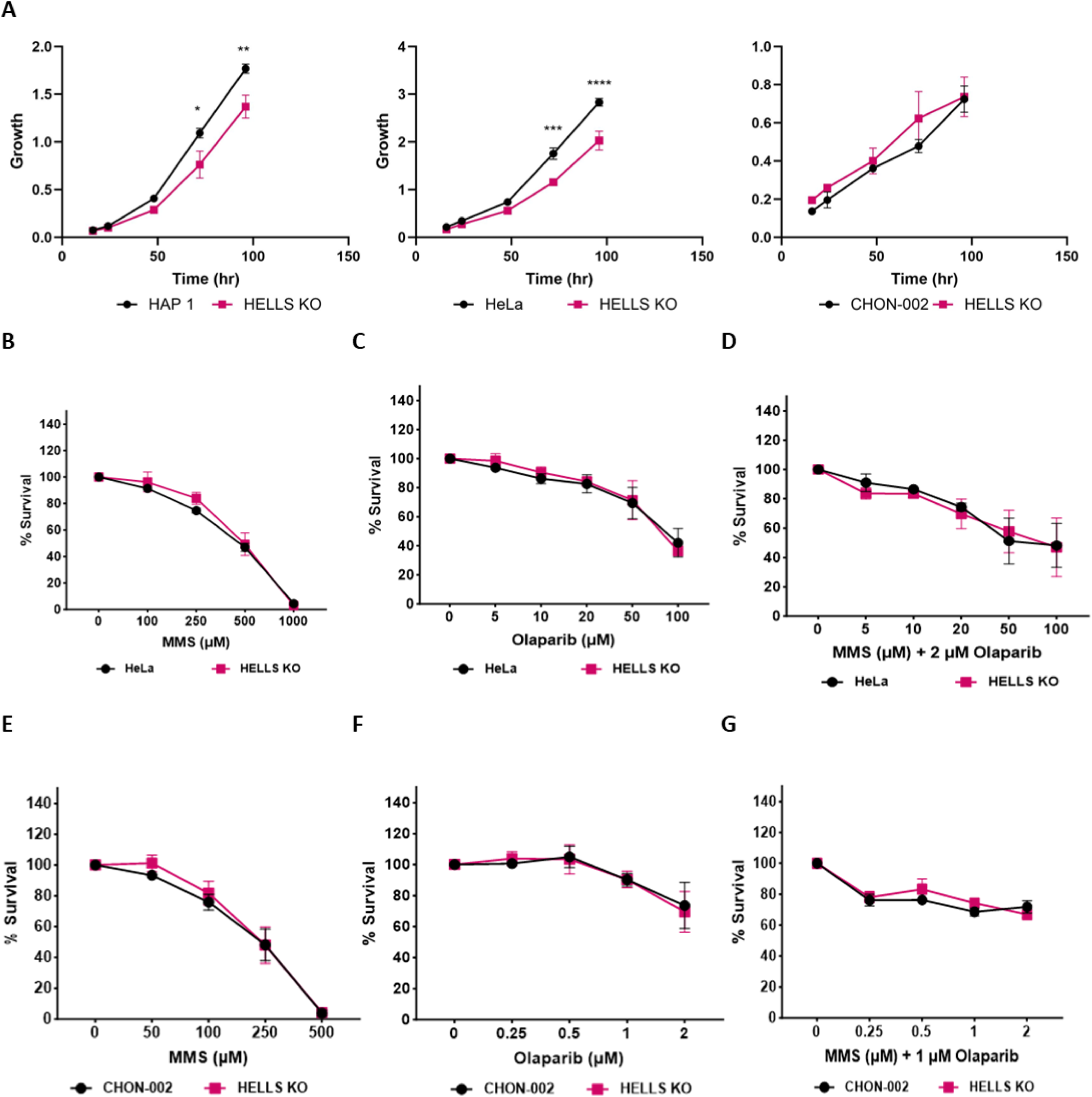
Loss of HELLS does not impact MMS and OLA sensitivity in HeLa and CHON-002 cells. **A.** Growth rate of parental and HELLS KO of HAPl, HeLa, and CHON-002 cell lines as measured by CCK-8 assay. Data are mean ± SEM. n=3 independent biological replicates. Statistical significance was determined by 2-way ANOVA using Sidak’s multiple comparison test. **B-D.** Cytotoxicity of HeLa parental and HELLS KO cells to indicated concentrations of **B.** MMS, C. Olaparib, and D. MMS and Olaparib for 48hr, tested using CCK-8 reagent. Data are mean ± SEM. n=3 independent biological replicates. E­ **G.** Cytotoxicity of CHON-002 parental and HELLS KO cells to indicated concentrations of E. MMS, F. Olaparib, and **G.** MMS and Olaparib for 48hr, tested using CCK-8 reagent. Data are mean ± SEM. n=3 independent biological replicates. (*p<0.05, **p<0.01, ***p<0.001, ****p<0.0001)

**Figure S2.**
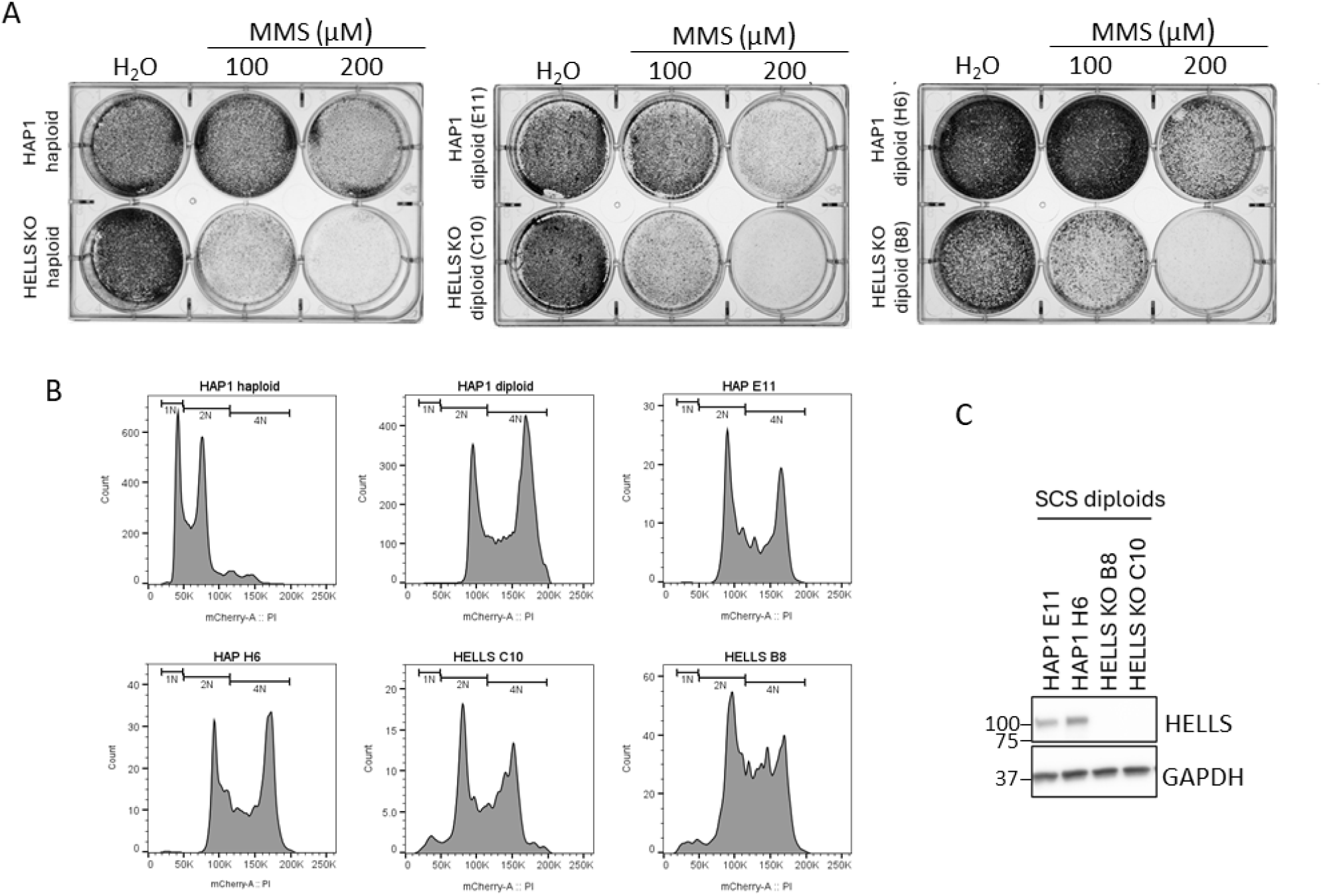
Loss of HELLS results in MMS sensitivity in both haploid and diploid clones of HAP1. **A.** Two independent single cell sorted diploid HAP1 and HELLS KO clones in a cell survival assay. **B.** Cell cycle histograms of single cell sorted cells from **A. C.** Western blot of single cells sorted clones from A. indicating HELLS expression level. GAPDH is used as a loading control.

**Figure S3.**
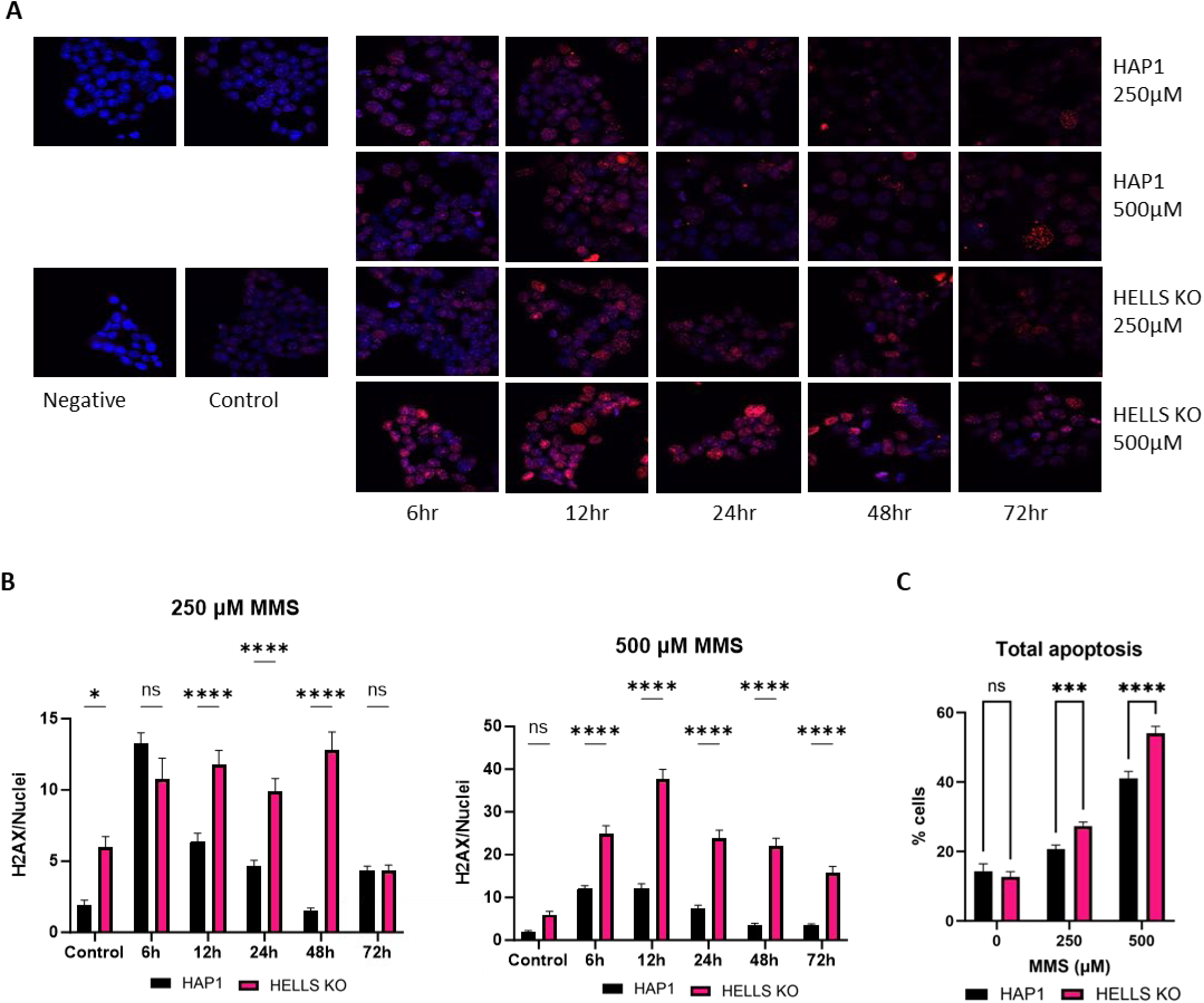
Loss of HELLS in HAPl leads to accumulation of alkylation-derived DNA damage and apoptosis. **A.** lmmunofluorescence-based detection and **B.** quantification of yH2AX in the nuclei of HELLS proficient and deficient cells. Cells were treated for 1hr at the indicated concentration of MMS before fixation. (n=9 pictures) Bonferroni test was used to determine significance. Data represents at least two independent experiments C. Proportion of apoptotic parental and HELLS KO HAP1 cells after treatment with MMS for 1hr. Data are mean ± SD. n=3 independent biological replicates. Two-way ANOVA using Tukey’s multiple comparisons test was used to determine significance. (*p<0.05, **p<0.01, ***p<0.001, ****p<0.0001)

**Figure S4.**
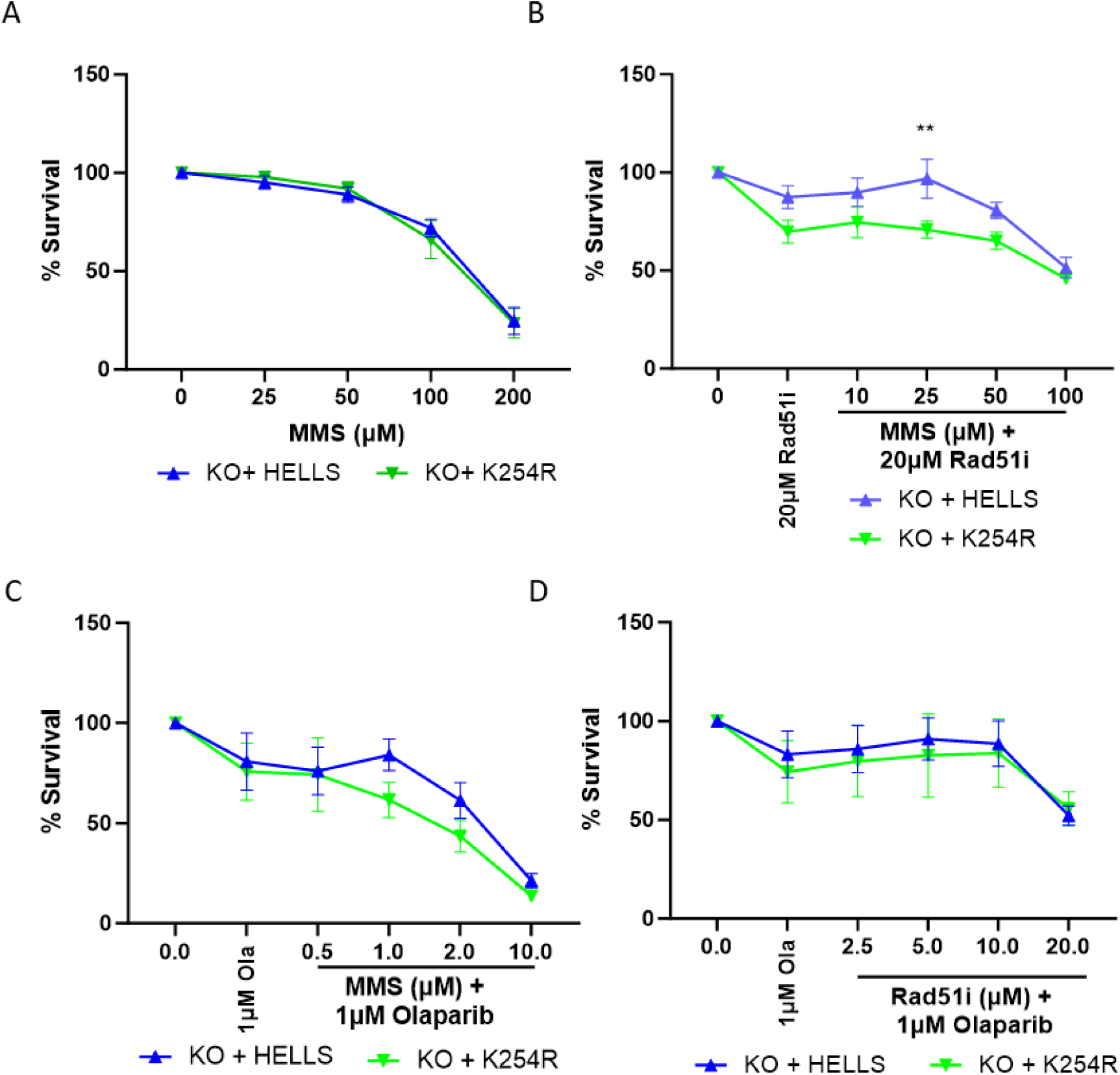
The ATPase function of the HELLS catalytic domain is not required for sensitivity to DNA damaging agents. CCK-8 survival assays for HAP-HELLS KO cells containing the WT HELLS gene or HELLS K254R after treatment for 48hr with **A.** MMS **B.** 20µM Rad51i and MMS **C.** 1µM Olaparib and MMS or D. 1µM Olaparib and Rad51i.

**Figure S5.**
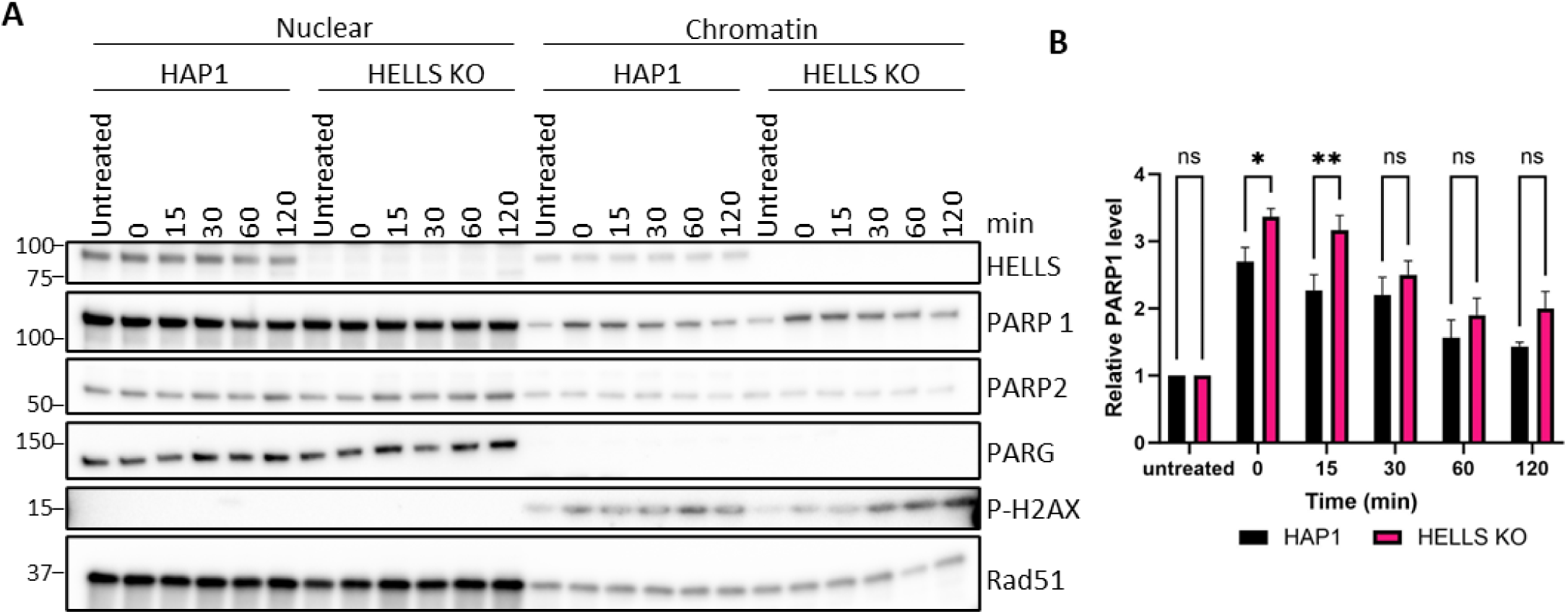
Loss of HELLS minimally impacts PARP1 trapping on chromatin. **A.** Representative western blot demonstrating levels of PARP1 captured on the chromatin. Cells were treated with 1mM MMS and 10µM Olaparib for 1hr and nuclear and chromatin fractions were resolved by SOS-PAGE and visualized by Western blot. **B.** Quantification of PARP1 levels of 3 biological replicates of **A.** Data are mean± SEM. Two-way ANOVA using Tukey’s multiple comparisons test was used to determine significance. (*p<0.05, **p<0.01)

**Figure S6.**
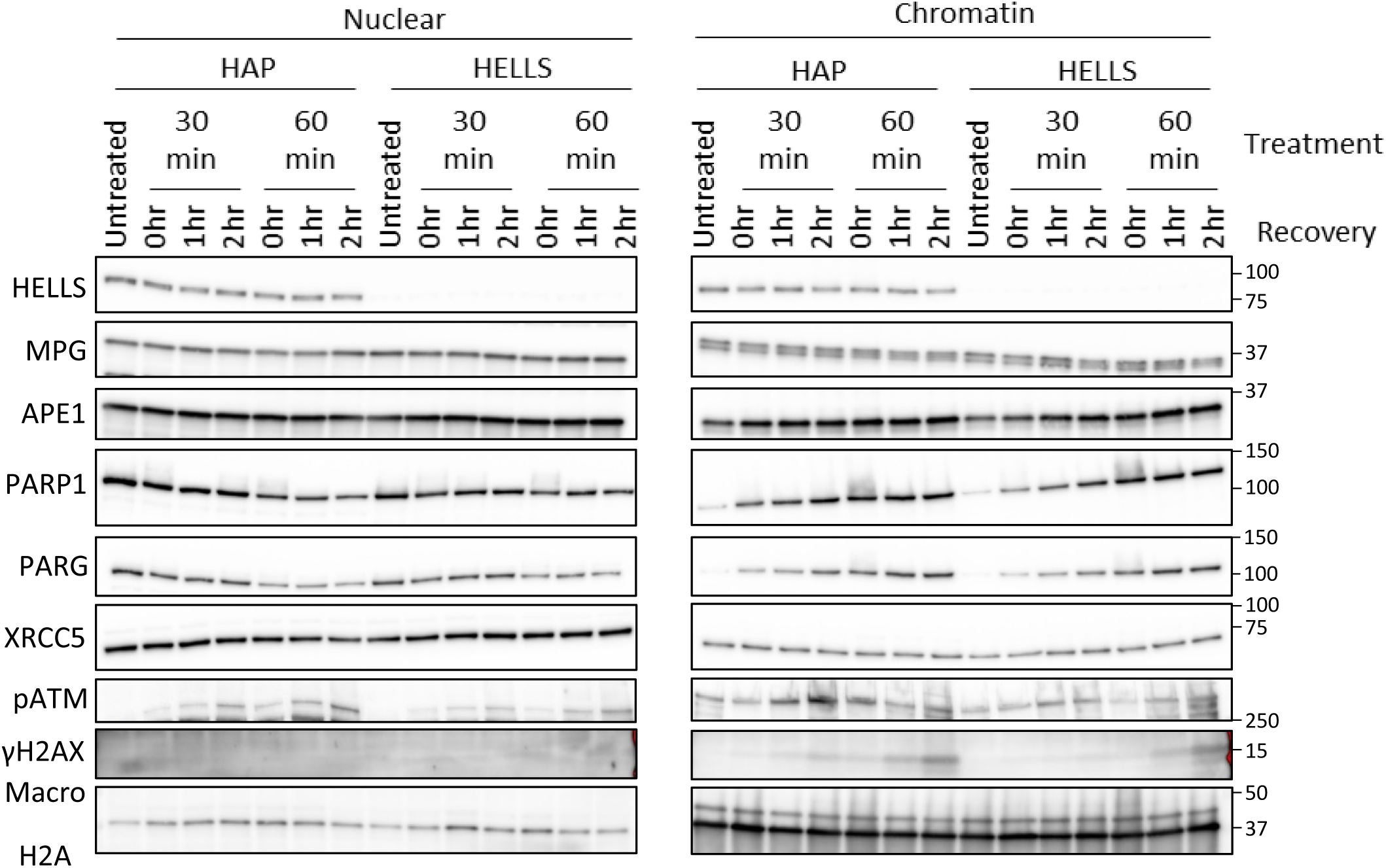
Loss of HELLS does not impact recruitment of select DNA repair proteins to chromatin in response to MMS. Cells were exposed to 3mM MMS for 30 and 60min, followed by drug removal and cell recovery in fresh media for 1 and 2hr. The nuclear and chromatin protein fractions were separated and detected by western blotting. Data represent at least two independent biological experiments.

## References

1. Caldecott, K. W., DNA single-strand break repair and human genetic disease. Trends Cell Biol 2022, 32, (9), 733–745.

2. SenGupta, T.; Palikaras, K.; Esbensen, Y. Q.; Konstantinidis, G.; Galindo, F. J. N.; Achanta, K.; Kassahun, H.; Stavgiannoudaki, I.; Bohr, V. A.; Akbari, M.; Gaare, J.; Tzoulis, C.; Tavernarakis, N.; Nilsen, H., Base excision repair causes age-dependent accumulation of single-stranded DNA breaks that contribute to Parkinson disease pathology. Cell Rep 2021, 36, (10), 109668.

3. Eustermann, S.; Patel, A. B.; Hopfner, K. P.; He, Y.; Korber, P., Energy-driven genome regulation by ATP-dependent chromatin remodellers. Nat Rev Mol Cell Biol 2024, 25, (4), 309–332.

4. Gourisankar, S.; Krokhotin, A.; Wenderski, W.; Crabtree, G. R., Context-specific functions of chromatin remodellers in development and disease. Nat Rev Genet 2024, 25, (5), 340–361.

5. Lans, H.; Marteijn, J. A.; Vermeulen, W., ATP-dependent chromatin remodeling in the DNA-damage response. Epigenetics Chromatin 2012, 5, 4.

6. Mayes, K.; Qiu, Z.; Alhazmi, A.; Landry, J. W., ATP-dependent chromatin remodeling complexes as novel targets for cancer therapy. Adv Cancer Res 2014, 121, 183–233.

7. Peixoto, E.; Khan, A.; Lewis, Z. A.; Contreras-Galindo, R.; Czaja, W., The Chromatin Remodeler HELLS: A New Regulator in DNA Repair, Genome Maintenance, and Cancer. Int J Mol Sci 2022, 23, (16).

8. Xiao, L.; Parolia, A.; Qiao, Y.; Bawa, P.; Eyunni, S.; Mannan, R.; Carson, S. E.; Chang, Y.; Wang, X.; Zhang, Y.; Vo, J. N.; Kregel, S.; Simko, S. A.; Delekta, A. D.; Jaber, M.; Zheng, H.; Apel, I. J.; McMurry, L.; Su, F.; Wang, R.; Zelenka-Wang, S.; Sasmal, S.; Khare, L.; Mukherjee, S.; Abbineni, C.; Aithal, K.; Bhakta, M. S.; Ghurye, J.; Cao, X.; Navone, N. M.; Nesvizhskii, A. I.; Mehra, R.; Vaishampayan, U.; Blanchette, M.; Wang, Y.; Samajdar, S.; Ramachandra, M.; Chinnaiyan, A. M., Targeting SWI/SNF ATPases in enhancer-addicted prostate cancer. Nature 2022, 601, (7893), 434–439.

9. Soll, J. M.; Sobol, R. W.; Mosammaparast, N., Regulation of DNA Alkylation Damage Repair: Lessons and Therapeutic Opportunities. Trends Biochem Sci 2017, 42, (3), 206–218.

10. Trivedi, R. N.; Almeida, K. H.; Fornsaglio, J. L.; Schamus, S.; Sobol, R. W., The role of base excision repair in the sensitivity and resistance to temozolomide-mediated cell death. Cancer Res 2005, 65, (14), 6394–400.

11. Fu, D.; Calvo, J. A.; Samson, L. D., Balancing repair and tolerance of DNA damage caused by alkylating agents. Nat Rev Cancer 2012, 12, (2), 104–20.

12. Kondo, N.; Takahashi, A.; Ono, K.; Ohnishi, T., DNA damage induced by alkylating agents and repair pathways. J Nucleic Acids 2010, 2010, 543531.

13. Heacock, M. L.; Stefanick, D. F.; Horton, J. K.; Wilson, S. H., Alkylation DNA damage in combination with PARP inhibition results in formation of S-phase-dependent double-strand breaks. DNA Repair (Amst) 2010, 9, (8), 929–36.

14. Horton, J. K.; Stefanick, D. F.; Prasad, R.; Gassman, N. R.; Kedar, P. S.; Wilson, S. H., Base excision repair defects invoke hypersensitivity to PARP inhibition. Mol Cancer Res 2014, 12, (8), 1128–39.

15. Hinz, J. M.; Czaja, W., Facilitation of base excision repair by chromatin remodeling. DNA Repair (Amst) 2015, 36, 91–97.

16. Meira, L. B.; Moroski-Erkul, C. A.; Green, S. L.; Calvo, J. A.; Bronson, R. T.; Shah, D.; Samson, L. D., Aag-initiated base excision repair drives alkylation-induced retinal degeneration in mice. Proc Natl Acad Sci U S A 2009, 106, (3), 888–93.

17. Rodriguez, Y.; Howard, M. J.; Cuneo, M. J.; Prasad, R.; Wilson, S. H., Unencumbered Pol beta lyase activity in nucleosome core particles. Nucleic Acids Res 2017, 45, (15), 8901–8915.

18. Tallis, M.; Morra, R.; Barkauskaite, E.; Ahel, I., Poly(ADP-ribosyl)ation in regulation of chromatin structure and the DNA damage response. Chromosoma 2014, 123, (1-2), 79–90.

19. Lin, X.; Leung, K. S. K.; Wolfe, K. F.; Call, N.; Bhandari, S. K.; Huang, X.; Lee, B. J.; Tomkinson, A. E.; Zha, S., XRCC1 mediates PARP1- and PAR-dependent recruitment of PARP2 to DNA damage sites. Nucleic Acids Res 2025, 53, (4).

20. Gohil, D.; Sarker, A. H.; Roy, R., Base Excision Repair: Mechanisms and Impact in Biology, Disease, and Medicine. Int J Mol Sci 2023, 24, (18).

21. Vrtis, K. B.; Dewar, J. M.; Chistol, G.; Wu, R. A.; Graham, T. G. W.; Walter, J. C., Single-strand DNA breaks cause replisome disassembly. Mol Cell 2021, 81, (6), 1309–1318 e6.

22. Hopkins, T. A.; Shi, Y.; Rodriguez, L. E.; Solomon, L. R.; Donawho, C. K.; DiGiammarino, E. L.; Panchal, S. C.; Wilsbacher, J. L.; Gao, W.; Olson, A. M.; Stolarik, D. F.; Osterling, D. J.; Johnson, E. F.; Maag, D., Mechanistic Dissection of PARP1 Trapping and the Impact on In Vivo Tolerability and Efficacy of PARP Inhibitors. Mol Cancer Res 2015, 13, (11), 1465–77.

23. Murai, J.; Huang, S. Y.; Das, B. B.; Renaud, A.; Zhang, Y.; Doroshow, J. H.; Ji, J.; Takeda, S.; Pommier, Y., Trapping of PARP1 and PARP2 by Clinical PARP Inhibitors. Cancer Res 2012, 72, (21), 5588–99.

24. Zhang, H.; Lin, X.; Zha, S., Revisiting PARP2 and PARP1 trapping through quantitative live-cell imaging. Biochem Soc Trans 2022, 50, (4), 1169–1177.

25. Hopkins, T. A.; Ainsworth, W. B.; Ellis, P. A.; Donawho, C. K.; DiGiammarino, E. L.; Panchal, S. C.; Abraham, V. C.; Algire, M. A.; Shi, Y.; Olson, A. M.; Johnson, E. F.; Wilsbacher, J. L.; Maag, D., PARP1 Trapping by PARP Inhibitors Drives Cytotoxicity in Both Cancer Cells and Healthy Bone Marrow. Mol Cancer Res 2019, 17, (2), 409–419.

26. Blessing, C.; Mandemaker, I. K.; Gonzalez-Leal, C.; Preisser, J.; Schomburg, A.; Ladurner, A. G., The Oncogenic Helicase ALC1 Regulates PARP Inhibitor Potency by Trapping PARP2 at DNA Breaks. Mol Cell 2020, 80, (5), 862–875 e6.

27. Juhasz, S.; Smith, R.; Schauer, T.; Spekhardt, D.; Mamar, H.; Zentout, S.; Chapuis, C.; Huet, S.; Timinszky, G., The chromatin remodeler ALC1 underlies resistance to PARP inhibitor treatment. Sci Adv 2020, 6, (51).

28. Krastev, D. B.; Li, S.; Sun, Y.; Wicks, A. J.; Hoslett, G.; Weekes, D.; Badder, L. M.; Knight, E. G.; Marlow, R.; Pardo, M. C.; Yu, L.; Talele, T. T.; Bartek, J.; Choudhary, J. S.; Pommier, Y.; Pettitt, S. J.; Tutt, A. N. J.; Ramadan, K.; Lord, C. J., The ubiquitin-dependent ATPase p97 removes cytotoxic trapped PARP1 from chromatin. Nat Cell Biol 2022, 24, (1), 62–73.

29. Demin, A. A.; Hirota, K.; Tsuda, M.; Adamowicz, M.; Hailstone, R.; Brazina, J.; Gittens, W.; Kalasova, I.; Shao, Z.; Zha, S.; Sasanuma, H.; Hanzlikova, H.; Takeda, S.; Caldecott, K. W., XRCC1 prevents toxic PARP1 trapping during DNA base excision repair. Mol Cell 2021, 81, (14), 3018–3030 e5.

30. Lee, D. W.; Zhang, K.; Ning, Z. Q.; Raabe, E. H.; Tintner, S.; Wieland, R.; Wilkins, B. J.; Kim, J. M.; Blough, R. I.; Arceci, R. J., Proliferation-associated SNF2-like gene (PASG): a SNF2 family member altered in leukemia. Cancer Res 2000, 60, (13), 3612–22.

31. Thijssen, P. E.; Ito, Y.; Grillo, G.; Wang, J.; Velasco, G.; Nitta, H.; Unoki, M.; Yoshihara, M.; Suyama, M.; Sun, Y.; Lemmers, R. J.; de Greef, J. C.; Gennery, A.; Picco, P.; Kloeckener-Gruissem, B.; Gungor, T.; Reisli, I.; Picard, C.; Kebaili, K.; Roquelaure, B.; Iwai, T.; Kondo, I.; Kubota, T.; van Ostaijen-Ten Dam, M. M.; van Tol, M. J.; Weemaes, C.; Francastel, C.; van der Maarel, S. M.; Sasaki, H., Mutations in CDCA7 and HELLS cause immunodeficiency-centromeric instability-facial anomalies syndrome. Nat Commun 2015, 6, 7870.

32. Liang, X.; Li, L.; Fan, Y., Diagnostic, Prognostic, and Immunological Roles of HELLS in Pan-Cancer: A Bioinformatics Analysis. Front Immunol 2022, 13, 870726.

33. Zhang, G.; Dong, Z.; Prager, B. C.; Kim, L. J.; Wu, Q.; Gimple, R. C.; Wang, X.; Bao, S.; Hamerlik, P.; Rich, J. N., Chromatin remodeler HELLS maintains glioma stem cells through E2F3 and MYC. JCI Insight 2019, 4, (7).

34. Zhu, W.; Li, L. L.; Songyang, Y.; Shi, Z.; Li, D., Identification and validation of HELLS (Helicase, Lymphoid-Specific) and ICAM1 (Intercellular adhesion molecule 1) as potential diagnostic biomarkers of lung cancer. PeerJ 2020, 8, e8731.

35. Burrage, J.; Termanis, A.; Geissner, A.; Myant, K.; Gordon, K.; Stancheva, I., The SNF2 family ATPase LSH promotes phosphorylation of H2AX and efficient repair of DNA double-strand breaks in mammalian cells. J Cell Sci 2012, 125, (Pt 22), 5524–34.

36. Kollarovic, G.; Topping, C. E.; Shaw, E. P.; Chambers, A. L., The human HELLS chromatin remodelling protein promotes end resection to facilitate homologous recombination and contributes to DSB repair within heterochromatin. Nucleic Acids Res 2020, 48, (4), 1872–1885.

37. Unoki, M.; Funabiki, H.; Velasco, G.; Francastel, C.; Sasaki, H., CDCA7 and HELLS mutations undermine nonhomologous end joining in centromeric instability syndrome. J Clin Invest 2019, 129, (1), 78–92.

38. Xu, X.; Ni, K.; He, Y.; Ren, J.; Sun, C.; Liu, Y.; Aladjem, M. I.; Burkett, S.; Finney, R.; Ding, X.; Sharan, S. K.; Muegge, K., The epigenetic regulator LSH maintains fork protection and genomic stability via MacroH2A deposition and RAD51 filament formation. Nat Commun 2021, 12, (1), 3520.

39. Liu, X.; Hou, X.; Zhou, Y.; Li, Q.; Kong, F.; Yan, S.; Lei, S.; Xiong, L.; He, J., Downregulation of the Helicase Lymphoid-Specific (HELLS) Gene Impairs Cell Proliferation and Induces Cell Cycle Arrest in Colorectal Cancer Cells. Onco Targets Ther 2019, 12, 10153–10163.

40. Basenko, E. Y.; Kamei, M.; Ji, L.; Schmitz, R. J.; Lewis, Z. A., The LSH/DDM1 Homolog MUS-30 Is Required for Genome Stability, but Not for DNA Methylation in Neurospora crassa. PLoS Genet 2016, 12, (1), e1005790.

41. Litwin, I.; Bakowski, T.; Maciaszczyk-Dziubinska, E.; Wysocki, R., The LSH/HELLS homolog Irc5 contributes to cohesin association with chromatin in yeast. Nucleic Acids Res 2017, 45, (11), 6404–6416.

42. Yao, Y.; Bilichak, A.; Golubov, A.; Kovalchuk, I., ddm1 plants are sensitive to methyl methane sulfonate and NaCl stresses and are deficient in DNA repair. Plant Cell Rep 2012, 31, (9), 1549–61.

43. Han, Y.; Ren, J.; Lee, E.; Xu, X.; Yu, W.; Muegge, K., Lsh/HELLS regulates self-renewal/proliferation of neural stem/progenitor cells. Sci Rep 2017, 7, (1), 1136.

44. Ren, J.; Finney, R.; Ni, K.; Cam, M.; Muegge, K., The chromatin remodeling protein Lsh alters nucleosome occupancy at putative enhancers and modulates binding of lineage specific transcription factors. Epigenetics 2019, 14, (3), 277–293.

45. Yan, Q.; Huang, J.; Fan, T.; Zhu, H.; Muegge, K., Lsh, a modulator of CpG methylation, is crucial for normal histone methylation. EMBO J 2003, 22, (19), 5154–62.

46. Boon, N. J.; Oliveira, R. A.; Korner, P. R.; Kochavi, A.; Mertens, S.; Malka, Y.; Voogd, R.; van der Horst, S. E. M.; Huismans, M. A.; Smabers, L. P.; Draper, J. M.; Wessels, L. F. A.; Haahr, P.; Roodhart, J. M. L.; Schumacher, T. N. M.; Snippert, H. J.; Agami, R.; Brummelkamp, T. R., DNA damage induces p53-independent apoptosis through ribosome stalling. Science 2024, 384, (6697), 785–792.

47. Dewan, A.; Xing, M.; Lundbaek, M. B.; Gago-Fuentes, R.; Beck, C.; Aas, P. A.; Liabakk, N. B.; Saeterstad, S.; Chau, K. T. P.; Kavli, B. M.; Oksenych, V., Robust DNA repair in PAXX-deficient mammalian cells. FEBS Open Bio 2018, 8, (3), 442–448.

48. Martins, M. B.; Perez, A. M.; Bohr, V. A.; Wilson, D. M., 3rd; Kobarg, J., NEK1 deficiency affects mitochondrial functions and the transcriptome of key DNA repair pathways. Mutagenesis 2021, 36, (3), 223–236.

49. Beigl, T. B.; Kjosas, I.; Seljeseth, E.; Glomnes, N.; Aksnes, H., Efficient and crucial quality control of HAP1 cell ploidy status. Biol Open 2020, 9, (11).

50. Li, X. C.; Tye, B. K., Ploidy dictates repair pathway choice under DNA replication stress. Genetics 2011, 187, (4), 1031–40.

51. Groth, P.; Auslander, S.; Majumder, M. M.; Schultz, N.; Johansson, F.; Petermann, E.; Helleday, T., Methylated DNA causes a physical block to replication forks independently of damage signalling, O(6)-methylguanine or DNA single-strand breaks and results in DNA damage. J Mol Biol 2010, 402, (1), 70–82.

52. Brem, R.; Hall, J., XRCC1 is required for DNA single-strand break repair in human cells. Nucleic Acids Res 2005, 33, (8), 2512–20.

53. Fenech, M.; Knasmueller, S.; Bolognesi, C.; Holland, N.; Bonassi, S.; Kirsch-Volders, M., Micronuclei as biomarkers of DNA damage, aneuploidy, inducers of chromosomal hypermutation and as sources of pro-inflammatory DNA in humans. Mutat Res Rev Mutat Res 2020, 786, 108342.

54. Lord, C. J.; Ashworth, A., The DNA damage response and cancer therapy. Nature 2012, 481, (7381), 287–94.

55. Lundin, C.; North, M.; Erixon, K.; Walters, K.; Jenssen, D.; Goldman, A. S.; Helleday, T., Methyl methanesulfonate (MMS) produces heat-labile DNA damage but no detectable in vivo DNA double-strand breaks. Nucleic Acids Res 2005, 33, (12), 3799–811.

56. Ensminger, M.; Iloff, L.; Ebel, C.; Nikolova, T.; Kaina, B.; Lbrich, M., DNA breaks and chromosomal aberrations arise when replication meets base excision repair. J Cell Biol 2014, 206, (1), 29–43.

57. Tameni, A.; Mallia, S.; Manicardi, V.; Donati, B.; Torricelli, F.; Vitale, E.; Salviato, E.; Gambarelli, G.; Muccioli, S.; Zanelli, M.; Ascani, S.; Martino, G.; Sanguedolce, F.; Sauta, E.; Tamagnini, I.; Puccio, N.; Neri, A.; Ciarrocchi, A.; Fragliasso, V., HELLS regulates transcription in T-cell lymphomas by reducing unscheduled R-loops and by facilitating RNAPII progression. Nucleic Acids Res 2024, 52, (11), 6171–6182.

58. Chou, W. C.; Wang, H. C.; Wong, F. H.; Ding, S. L.; Wu, P. E.; Shieh, S. Y.; Shen, C. Y., Chk2-dependent phosphorylation of XRCC1 in the DNA damage response promotes base excision repair. EMBO J 2008, 27, (23), 3140–50.

59. Ronson, G. E.; Piberger, A. L.; Higgs, M. R.; Olsen, A. L.; Stewart, G. S.; McHugh, P. J.; Petermann, E.; Lakin, N. D., PARP1 and PARP2 stabilise replication forks at base excision repair intermediates through Fbh1-dependent Rad51 regulation. Nat Commun 2018, 9, (1), 746.

60. Bonilla, B.; Hengel, S. R.; Grundy, M. K.; Bernstein, K. A., RAD51 Gene Family Structure and Function. Annu Rev Genet 2020, 54, 25–46.

61. Gonzalez-Prieto, R.; Munoz-Cabello, A. M.; Cabello-Lobato, M. J.; Prado, F., Rad51 replication fork recruitment is required for DNA damage tolerance. EMBO J 2013, 32, (9), 1307–21.

62. Hanzlikova, H.; Caldecott, K. W., Perspectives on PARPs in S Phase. Trends Genet 2019, 35, (6), 412–422.

63. Rose, M.; Burgess, J. T.; O’Byrne, K.; Richard, D. J.; Bolderson, E., PARP Inhibitors: Clinical Relevance, Mechanisms of Action and Tumor Resistance. Front Cell Dev Biol 2020, 8, 564601.

64. Bajrami, I.; Frankum, J. R.; Konde, A.; Miller, R. E.; Rehman, F. L.; Brough, R.; Campbell, J.; Sims, D.; Rafiq, R.; Hooper, S.; Chen, L.; Kozarewa, I.; Assiotis, I.; Fenwick, K.; Natrajan, R.; Lord, C. J.; Ashworth, A., Genome-wide profiling of genetic synthetic lethality identifies CDK12 as a novel determinant of PARP1/2 inhibitor sensitivity. Cancer Res 2014, 74, (1), 287–97.

65. Hassan, S.; Esch, A.; Liby, T.; Gray, J. W.; Heiser, L. M., Pathway-Enriched Gene Signature Associated with 53BP1 Response to PARP Inhibition in Triple-Negative Breast Cancer. Mol Cancer Ther 2017, 16, (12), 2892–2901.

66. Tsujino, T.; Takai, T.; Hinohara, K.; Gui, F.; Tsutsumi, T.; Bai, X.; Miao, C.; Feng, C.; Gui, B.; Sztupinszki, Z.; Simoneau, A.; Xie, N.; Fazli, L.; Dong, X.; Azuma, H.; Choudhury, A. D.; Mouw, K. W.; Szallasi, Z.; Zou, L.; Kibel, A. S.; Jia, L., CRISPR screens reveal genetic determinants of PARP inhibitor sensitivity and resistance in prostate cancer. Nat Commun 2023, 14, (1), 252.

67. Cong, K.; Peng, M.; Kousholt, A. N.; Lee, W. T. C.; Lee, S.; Nayak, S.; Krais, J.; VanderVere-Carozza, P. S.; Pawelczak, K. S.; Calvo, J.; Panzarino, N. J.; Turchi, J. J.; Johnson, N.; Jonkers, J.; Rothenberg, E.; Cantor, S. B., Replication gaps are a key determinant of PARP inhibitor synthetic lethality with BRCA deficiency. Mol Cell 2021, 81, (15), 3227.

68. Hewitt, G.; Borel, V.; Segura-Bayona, S.; Takaki, T.; Ruis, P.; Bellelli, R.; Lehmann, L. C.; Sommerova, L.; Vancevska, A.; Tomas-Loba, A.; Zhu, K.; Cooper, C.; Fugger, K.; Patel, H.; Goldstone, R.; Schneider-Luftman, D.; Herbert, E.; Stamp, G.; Brough, R.; Pettitt, S.; Lord, C. J.; West, S. C.; Ahel, I.; Ahel, D.; Chapman, J. R.; Deindl, S.; Boulton, S. J., Defective ALC1 nucleosome remodeling confers PARPi sensitization and synthetic lethality with HRD. Mol Cell 2021, 81, (4), 767–783 e11.

69. Ray Chaudhuri, A.; Nussenzweig, A., The multifaceted roles of PARP1 in DNA repair and chromatin remodelling. Nat Rev Mol Cell Biol 2017, 18, (10), 610–621.

70. Houl, J. H.; Ye, Z.; Brosey, C. A.; Balapiti-Modarage, L. P. F.; Namjoshi, S.; Bacolla, A.; Laverty, D.; Walker, B. L.; Pourfarjam, Y.; Warden, L. S.; Babu Chinnam, N.; Moiani, D.; Stegeman, R. A.; Chen, M. K.; Hung, M. C.; Nagel, Z. D.; Ellenberger, T.; Kim, I. K.; Jones, D. E.; Ahmed, Z.; Tainer, J. A., Selective small molecule PARG inhibitor causes replication fork stalling and cancer cell death. Nat Commun 2019, 10, (1), 5654.

71. Min, W.; Wang, Z. Q., Poly (ADP-ribose) glycohydrolase (PARG) and its therapeutic potential. Front Biosci (Landmark Ed) 2009, 14, (5), 1619–26.

72. Huang, F.; Mazin, A. V., A small molecule inhibitor of human RAD51 potentiates breast cancer cell killing by therapeutic agents in mouse xenografts. PLoS One 2014, 9, (6), e100993.

73. Wera, A. C.; Lobbens, A.; Stoyanov, M.; Lucas, S.; Michiels, C., Radiation-induced synthetic lethality: combination of poly(ADP-ribose) polymerase and RAD51 inhibitors to sensitize cells to proton irradiation. Cell Cycle 2019, 18, (15), 1770–1783.

74. Du, W.; Liu, X.; Yang, M.; Wang, W.; Sun, J., The Regulatory Role of PRRX1 in Cancer Epithelial-Mesenchymal Transition. Onco Targets Ther 2021, 14, 4223–4229.

75. Takihira, S.; Yamada, D.; Osone, T.; Takao, T.; Sakaguchi, M.; Hakozaki, M.; Itano, T.; Nakata, E.; Fujiwara, T.; Kunisada, T.; Ozaki, T.; Takarada, T., PRRX1-TOP2A interaction is a malignancy-promoting factor in human malignant peripheral nerve sheath tumours. Br J Cancer 2024, 130, (9), 1493–1504.

76. Centore, R. C.; Sandoval, G. J.; Soares, L. M. M.; Kadoch, C.; Chan, H. M., Mammalian SWI/SNF Chromatin Remodeling Complexes: Emerging Mechanisms and Therapeutic Strategies. Trends Genet 2020, 36, (12), 936–950.

77. Verma, P.; Zhou, Y.; Cao, Z.; Deraska, P. V.; Deb, M.; Arai, E.; Li, W.; Shao, Y.; Puentes, L.; Li, Y.; Patankar, S.; Mach, R. H.; Faryabi, R. B.; Shi, J.; Greenberg, R. A., ALC1 links chromatin accessibility to PARP inhibitor response in homologous recombination-deficient cells. Nat Cell Biol 2021, 23, (2), 160–171.

78. Caldecott, K. W., Causes and consequences of DNA single-strand breaks. Trends Biochem Sci 2024, 49, (1), 68–78.

79. Srinivas, U. S.; Tan, B. W. Q.; Vellayappan, B. A.; Jeyasekharan, A. D., ROS and the DNA damage response in cancer. Redox Biol 2019, 25, 101084.

80. Tell, G.; Demple, B., Base excision DNA repair and cancer. Oncotarget 2015, 6, (2), 584–5.

81. Wallace, S. S.; Murphy, D. L.; Sweasy, J. B., Base excision repair and cancer. *Cancer Lett* 2012, 327, (1–2), 73-89.

82. Ruiz, P. D.; Hamilton, G. A.; Park, J. W.; Gamble, M. J., MacroH2A1 Regulation of Poly(ADP-Ribose) Synthesis and Stability Prevents Necrosis and Promotes DNA Repair. Mol Cell Biol 2019, 40, (1).

83. Sebastian, R.; Hosogane, E. K.; Sun, E. G.; Tran, A. D.; Reinhold, W. C.; Burkett, S.; Sturgill, D. M.; Gudla, P. R.; Pommier, Y.; Aladjem, M. I.; Oberdoerffer, P., Epigenetic Regulation of DNA Repair Pathway Choice by MacroH2A1 Splice Variants Ensures Genome Stability. Mol Cell 2020, 79, (5), 836–845 e7.

84. Ni, K.; Muegge, K., LSH catalyzes ATP-driven exchange of histone variants macroH2A1 and macroH2A2. Nucleic Acids Res 2021, 49, (14), 8024–8036.

85. Sun, L. Q.; Lee, D. W.; Zhang, Q.; Xiao, W.; Raabe, E. H.; Meeker, A.; Miao, D.; Huso, D. L.; Arceci, R. J., Growth retardation and premature aging phenotypes in mice with disruption of the SNF2-like gene, PASG. Genes Dev 2004, 18, (9), 1035–46.

86. Tsuda, M.; Cho, K.; Ooka, M.; Shimizu, N.; Watanabe, R.; Yasui, A.; Nakazawa, Y.; Ogi, T.; Harada, H.; Agama, K.; Nakamura, J.; Asada, R.; Fujiike, H.; Sakuma, T.; Yamamoto, T.; Murai, J.; Hiraoka, M.; Koike, K.; Pommier, Y.; Takeda, S.; Hirota, K., ALC1/CHD1L, a chromatin-remodeling enzyme, is required for efficient base excision repair. PLoS One 2017, 12, (11), e0188320.

87. Serrano-Benitez, A.; Wells, S. E.; Drummond-Clarke, L.; Russo, L. C.; Thomas, J. C.; Leal, G. A.; Farrow, M.; Edgerton, J. M.; Balasubramanian, S.; Yang, M.; Frezza, C.; Gautam, A.; Brazina, J.; Burdova, K.; Hoch, N. C.; Jackson, S. P.; Caldecott, K. W., Unrepaired base excision repair intermediates in template DNA strands trigger replication fork collapse and PARP inhibitor sensitivity. EMBO J 2023, 42, (18), e113190.

88. Gill, S. J.; Travers, J.; Pshenichnaya, I.; Kogera, F. A.; Barthorpe, S.; Mironenko, T.; Richardson, L.; Benes, C. H.; Stratton, M. R.; McDermott, U.; Jackson, S. P.; Garnett, M. J., Combinations of PARP Inhibitors with Temozolomide Drive PARP1 Trapping and Apoptosis in Ewing’s Sarcoma. PLoS One 2015, 10, (10), e0140988.

89. Mahalingam, P.; Smith, S.; Lopez, J.; Sharma, R. K.; Millard, T.; Thway, K.; Fisher, C.; Reardon, D. A.; Jones, R.; Nicholson, A. G.; Cunningham, D.; Welsh, L.; Sharma, B., PARP inhibition utilized in combination therapy with Olaparib-Temozolomide to achieve disease stabilization in a rare case of BRCA1-mutant, metastatic myxopapillary ependymoma. Rare Tumors 2023, 15, 20363613231152333.

90. Zhang, J.; Gao, Y.; Zhang, Z.; Zhao, J.; Jia, W.; Xia, C.; Wang, F.; Liu, T., Multi-therapies Based on PARP Inhibition: Potential Therapeutic Approaches for Cancer Treatment. J Med Chem 2022, 65, (24), 16099–16127.

91. Kedar, P. S.; Stefanick, D. F.; Horton, J. K.; Wilson, S. H., Increased PARP-1 association with DNA in alkylation damaged, PARP-inhibited mouse fibroblasts. Mol Cancer Res 2012, 10, (3), 360–8.

92. Chai, W. X.; Sun, L. G.; Dai, F. H.; Shao, H. S.; Zheng, N. G.; Cai, H. Y., Inhibition of PRRX2 suppressed colon cancer liver metastasis via inactivation of Wnt/beta-catenin signaling pathway. Pathol Res Pract 2019, 215, (10), 152593.

93. Liu, L.; Liu, A.; Liu, X., PRRX2 predicts poor survival prognosis, and promotes malignant phenotype of lung adenocarcinoma via transcriptional activates PSMD1. Transl Oncol 2023, 27, 101586.

94. Lv, Z. D.; Wang, H. B.; Liu, X. P.; Jin, L. Y.; Shen, R. W.; Wang, X. G.; Kong, B.; Qu, H. L.; Li, F. N.; Yang, Q. F., Silencing of Prrx2 Inhibits the Invasion and Metastasis of Breast Cancer both In Vitro and In Vivo by Reversing Epithelial-Mesenchymal Transition. Cell Physiol Biochem 2017, 42, (5), 1847–1856.

95. Chen, Z.; Chen, Y.; Li, Y.; Lian, W.; Zheng, K.; Zhang, Y.; Zhang, Y.; Lin, C.; Liu, C.; Sun, F.; Sun, X.; Wang, J.; Zhao, L.; Ke, Y., Prrx1 promotes stemness and angiogenesis via activating TGF-beta/smad pathway and upregulating proangiogenic factors in glioma. Cell Death Dis 2021, 12, (6), 615.

96. Yao, J.; Zhang, Y.; Xia, Y.; Zhu, C.; Wen, X.; Liu, T.; Da, M., PRRX1 promotes lymph node metastasis of gastric cancer by regulating epithelial-mesenchymal transition. Medicine (Baltimore) 2021, 100, (6), e24674.

97. Moller, P.; Azqueta, A.; Boutet-Robinet, E.; Koppen, G.; Bonassi, S.; Milic, M.; Gajski, G.; Costa, S.; Teixeira, J. P.; Costa Pereira, C.; Dusinska, M.; Godschalk, R.; Brunborg, G.; Gutzkow, K. B.; Giovannelli, L.; Cooke, M. S.; Richling, E.; Laffon, B.; Valdiglesias, V.; Basaran, N.; Del Bo, C.; Zegura, B.; Novak, M.; Stopper, H.; Vodicka, P.; Vodenkova, S.; de Andrade, V. M.; Sramkova, M.; Gabelova, A.; Collins, A.; Langie, S. A. S., Minimum Information for Reporting on the Comet Assay (MIRCA): recommendations for describing comet assay procedures and results. Nat Protoc 2020, 15, (12), 3817–3826.

