## Supplementary data for "Chromatin regulator HELLS mediates SSB repair and responses to DNA alkylation damage"

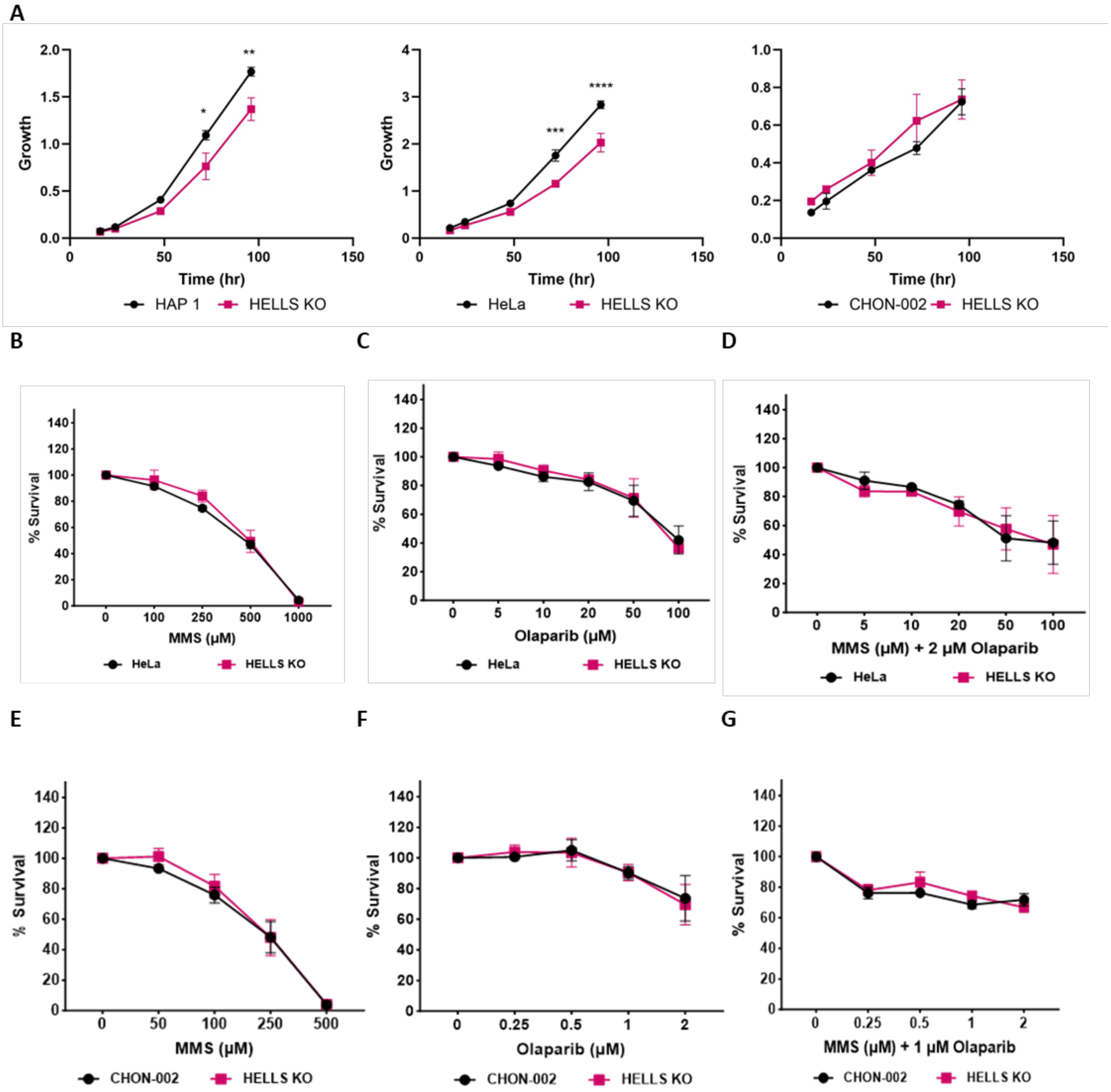


**Figure S1. Loss of HELLS does not impact MMS and OLA sensitivity in HeLa and CHON-002 cells. A.** Growth rate of parental and HELLS KO of HAP1, HeLa, and CHON-002 cell lines as measured by CCK-8 assay. Data are mean ± SEM. n=3 independent biological replicates. Statistical significance was determined by 2-way ANOVA using Sidak's multiple comparison test. **B-D.** Cytotoxicity of HeLa parental and HELLS KO cells to indicated concentrations of **B.** MMS, **C.** Olaparib, and **D.** MMS and Olaparib for 48hr, tested using CCK-8 reagent. Data are mean ± SEM. n=3 independent biological replicates. **E-G.** Cytotoxicity of CHON-002 parental and HELLS KO cells to indicated concentrations of **E.** MMS, **F.** Olaparib, and **G.** MMS and Olaparib for 48hr, tested using CCK-8 reagent. Data are mean ± SEM. n=3 independent biological replicates. (*p<0.05, **p<0.01, ***p<0.001, ****p<0.0001)

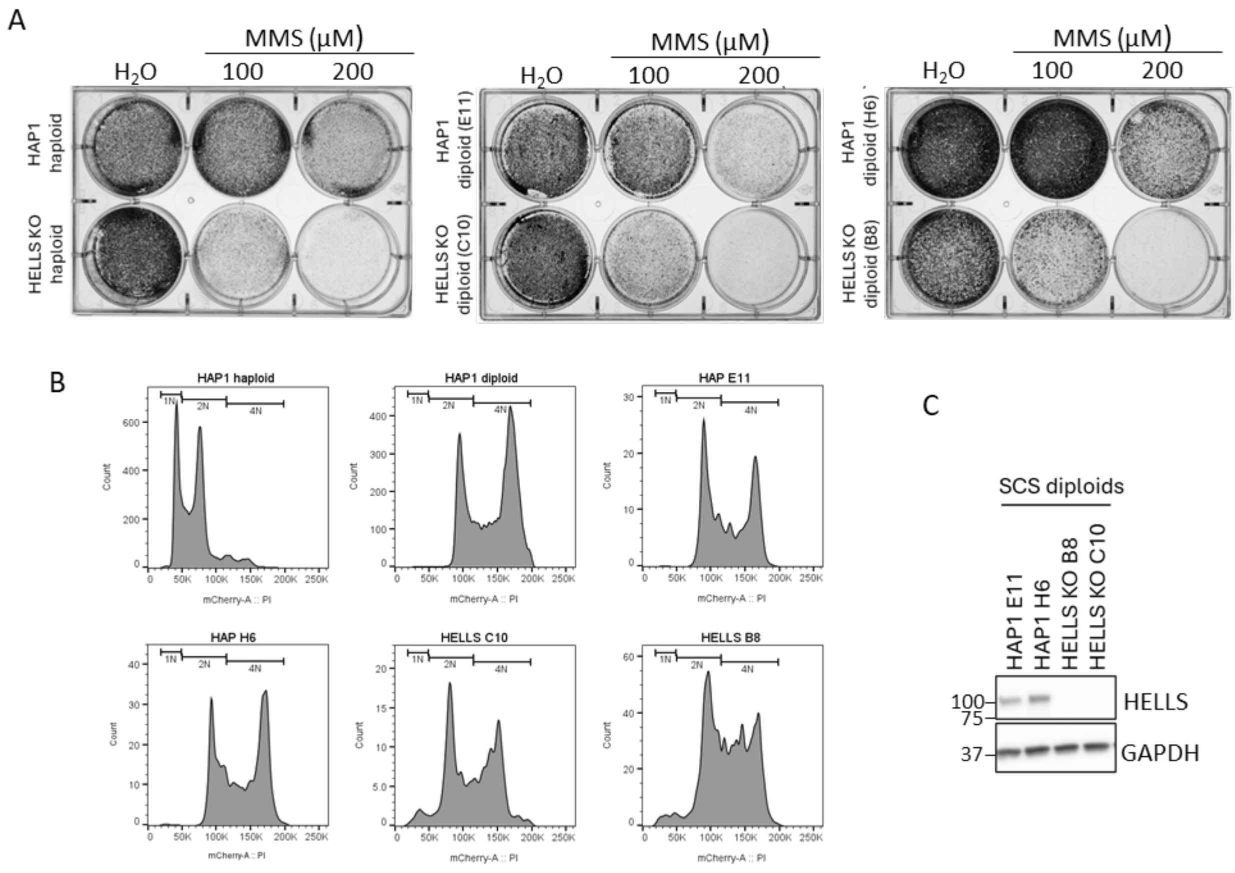


**Figure S2. Loss of HELLS results in MMS sensitivity in both haploid and diploid clones of HAP1. A.** Two independent single cell sorted diploid HAP1 and HELLS KO clones in a cell survival assay. **B.** Cell cycle histograms of single cell sorted cells from **A. C.** Western blot of single cells sorted clones from **A.** indicating HELLS expression level. GAPDH is used as a loading control.

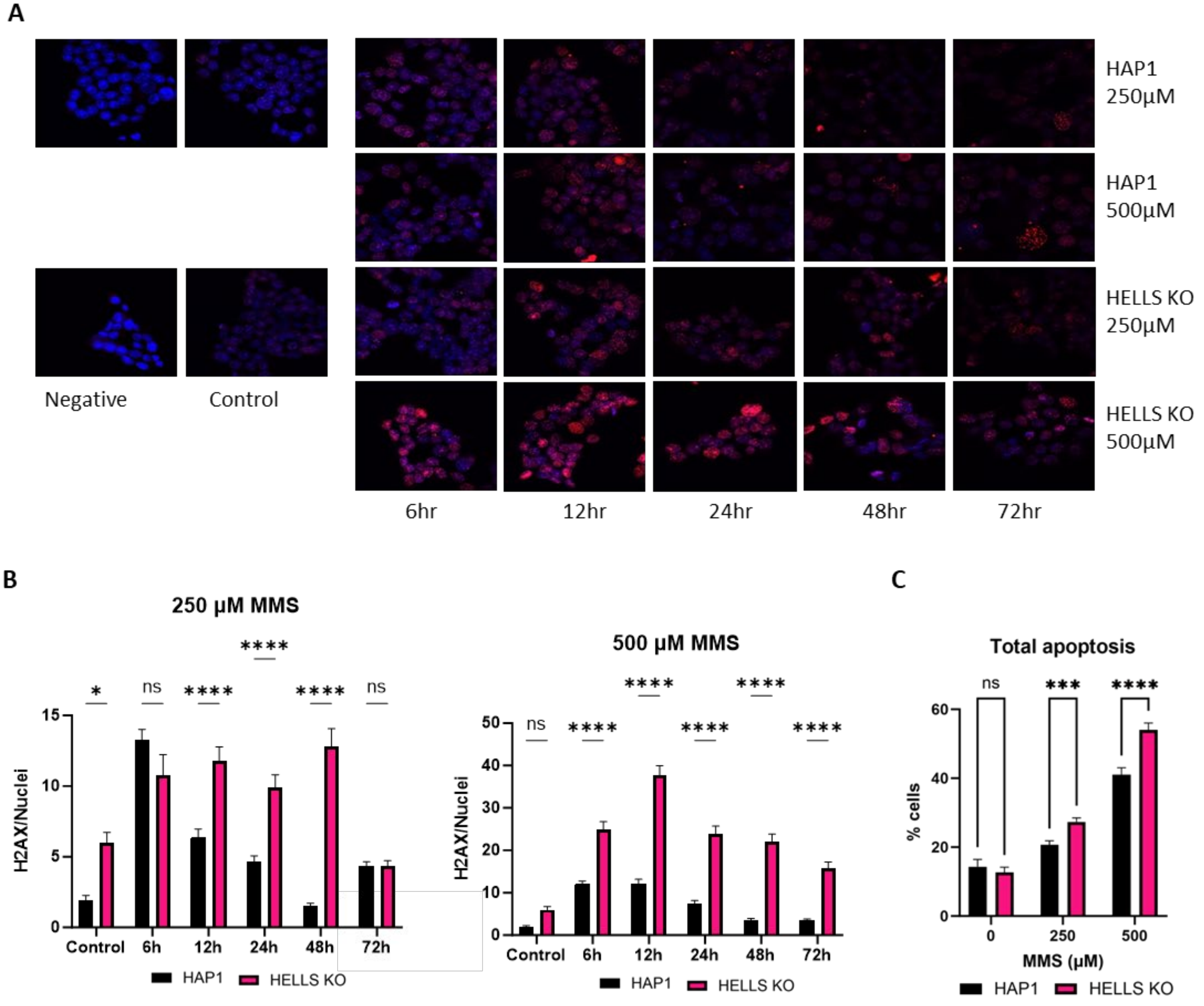


**Figure S3. Loss of HELLS in HAP1 leads to accumulation of alkylation-derived DNA damage and apoptosis. A.** Immunofluorescence-based detection and **B.** quantification of γH2AX in the nuclei of HELLS proficient and deficient cells. Cells were treated for 1hr at the indicated concentration of MMS before fixation. (n=9 pictures) Bonferroni test was used to determine significance. Data represents at least two independent experiments **C.** Proportion of apoptotic parental and HELLS KO HAP1 cells after treatment with MMS for 1hr. Data are mean ± SD. n=3 independent biological replicates. Two-way ANOVA using Tukey's multiple comparisons test was used to determine significance. (*p<0.05, **p<0.01, ***p<0.001, ****p<0.0001)

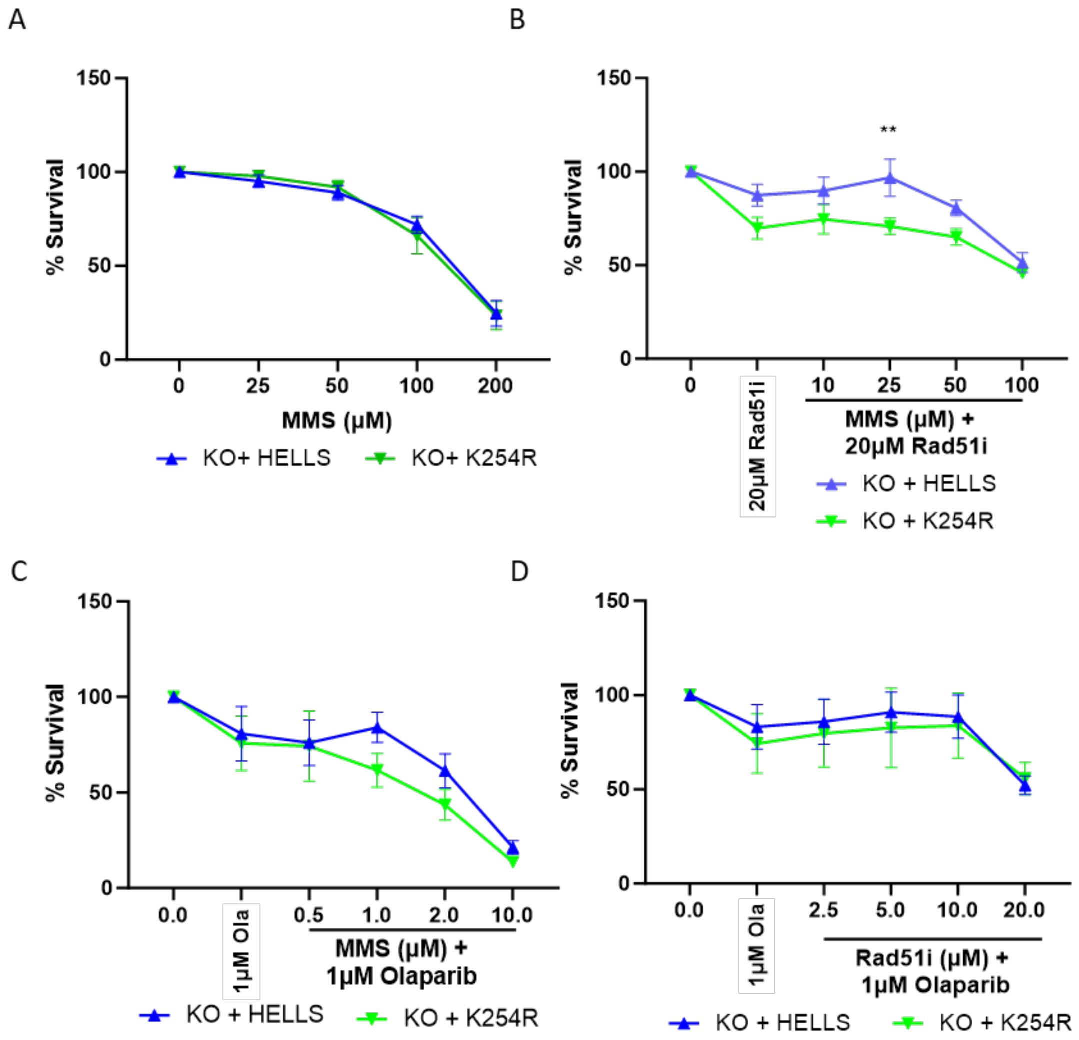


**Figure S4. The ATPase function of the HELLS catalytic domain is not required for sensitivity to DNA damaging agents.** CCK-8 survival assays for HAP-HELLS KO cells containing the WT HELLS gene or HELLS K254R after treatment for 48hr with **A**. MMS **B**. 20µM Rad51i and MMS **C**. 1µM Olaparib and MMS or **D**. 1µM Olaparib and Rad51i.

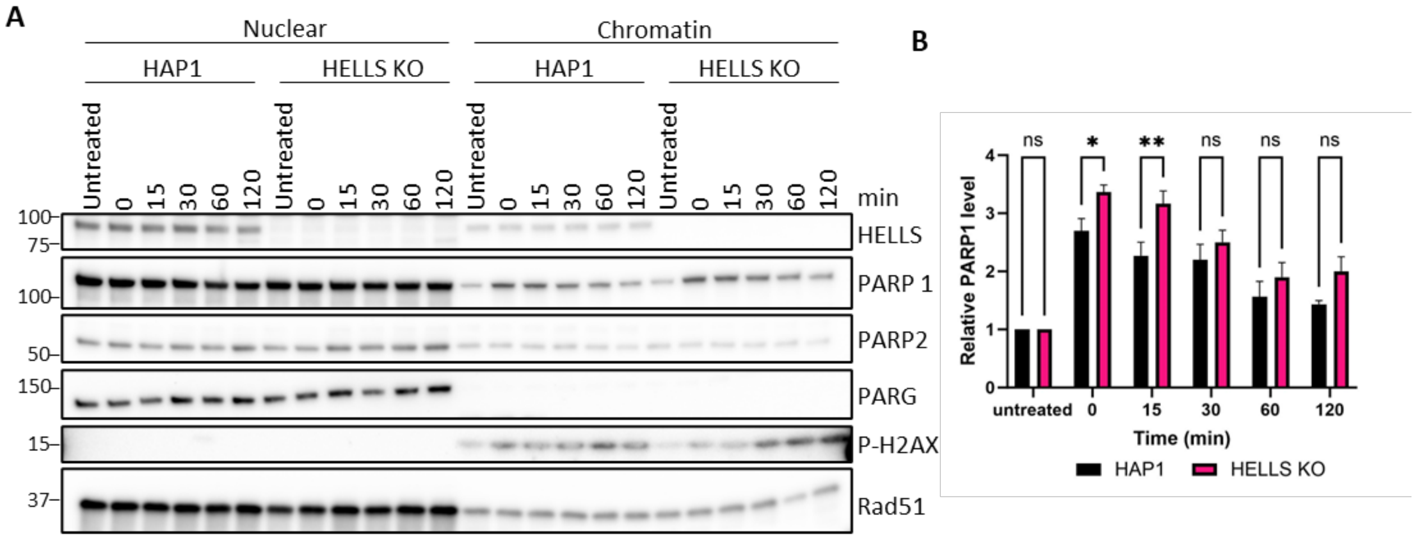


**Figure S5. Loss of HELLS minimally impacts PARP1 trapping on chromatin. A.** Representative western blot demonstrating levels of PARP1 captured on the chromatin. Cells were treated with 1mM MMS and 10µM Olaparib for 1hr and nuclear and chromatin fractions were resolved by SDS-PAGE and visualized by Western blot. **B.** Quantification of PARP1 levels of 3 biological replicates of **A.** Data are mean ± SEM. Two-way ANOVA using Tukey's multiple comparisons test was used to determine significance. (*$p<0.05$, **$p<0.01$)

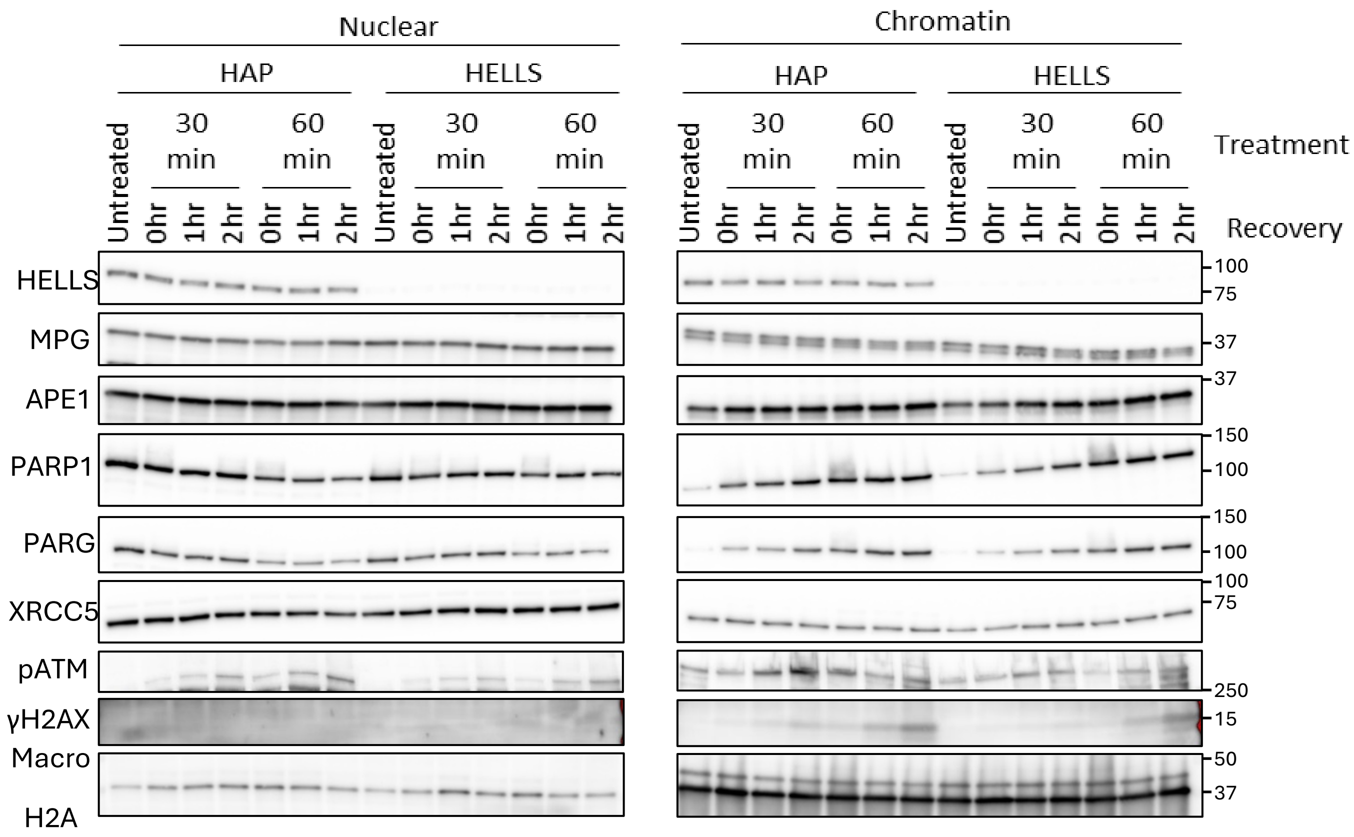


**Figure S6. Loss of HELLS does not impact recruitment of select DNA repair proteins to chromatin in response to MMS.** Cells were exposed to 3mM MMS for 30 and 60min, followed by drug removal and cell recovery in fresh media for 1 and 2hr. The nuclear and chromatin protein fractions were separated and detected by western blotting. Data represent at least two independent biological experiments.
